# A cross model spatial and single-cell atlas reveals the conserved involvement of osteopontin in polycystic kidney disease

**DOI:** 10.1101/2025.09.17.676846

**Authors:** Sarah J. Miller, Hua Zhong, Weidong Wu, Audrey M. Cordova, Morgan E. Smith, Alex Yashchenko, Zhang Li, Daniyal J. Jafree, Chelsea N. Zimmerman, Christa I. DeVette, Vicki Do, Maya Hignite, Isabella G. Darby, Khalecha Bintha Ahmed, Yohan Park, Fariha Nusrat, Bibi Maryam, Sizhao Lu, Xiaoyan Li, Jenny R. Gipson, Xiaogang Li, David A. Long, Mary Weiser-Evans, Bradley K. Yoder, Benjamin D. Cowley, Katharina Hopp, Jason Stubbs, Qin Ma, Anjun Ma, Kurt A. Zimmerman

**Author notes:** Corresponding Author: Kurt A. Zimmerman, Section of Nephrology, Department of Internal Medicine. University of Oklahoma Health Sciences Center Biomedical Sciences building, Room 513, Oklahoma City, Oklahoma 73104. authors contributed equally to this work (Sarah J. Miller, Hua Zhong).

## Abstract

Polycystic kidney disease (PKD) arises from mutations in cilia-associated genes, such as *PKD1* and *PKD2*, expressed in renal epithelial cells, leading to progressive kidney dysfunction and end-stage kidney disease (ESKD). PKD patients exhibit significant heterogeneity in disease progression, largely due to genetic and environmental modifiers. Like patients, mouse models of PKD also exhibit significant heterogeneity with regards to the gene mutated, age of disease onset, and rate of disease progression. To elucidate the cellular and molecular consequences of these variables, we constructed an integrated single-cell and spatial transcriptomics atlas across mouse models of PKD, mapping changes in cell type composition, gene expression, and intercellular signaling networks across the whole atlas and within individual models. Consistently across models, single cell RNA sequencing (scRNAseq) data revealed increased *Spp1* (osteopontin) expression and signaling from PKD-enriched clusters to Ly6c^lo^ monocytes. Global deletion of *Spp1* in *Pkd1*^RC/RC^ mice resulted in reduced cyst severity, improved kidney function, and reduced Ly6c^lo^ monocyte numbers, suggesting that SPP1 signaling to Ly6c^lo^ monocytes promotes PKD progression. We also created a freely available, searchable website (https://bmblx.bmi.osumc.edu/scPKD/) that can be used to identify cross- and intra-model specific changes in gene expression, guiding researchers to new therapeutic targets for treating PKD.

## Introduction

Autosomal dominant polycystic kidney disease (ADPKD) is caused by mutations in cilia-related genes including *PKD1* and *PKD2,* among others, and results in the development and growth of fluid-filled cysts throughout the kidney, eventually leading to end-stage kidney disease (ESKD)^1–3^. ADPKD patients have significant heterogeneity in the rate of disease progression, largely owing to the spectrum of genetic variants and environmental modifiers^4^. Like patients, mouse models of PKD also exhibit significant heterogeneity with regards to the gene mutated, age of disease onset, and rate of disease progression. Other differences between models include promoters that drive Cre recombinase activity, which can range from nephron-specific (*Pax8, Ksp*, or *Pkhd1*) to global (Cagg). Further adding to the complexity is the epistatic relationship between PKD genes (*Pkd1, Pkd2*) and primary cilia, as mice lacking both primary cilia and PKD-associated genes have less severe disease compared to mice lacking only PKD-associated genes^5^. Collectively, the large variation in PKD models significantly impairs our understanding of shared and model-specific cellular and molecular features of PKD.

In this manuscript, we generated a cross-model, spatial and single-cell RNA sequencing (scRNAseq) atlas to understand how PKD-associated mutations impact the abundance of kidney cell types and their molecular profiles across and within individual PKD models. Using our integrated atlas, we identified cross-model dysregulation of osteopontin (SPP1) signaling and validate this observation by demonstrating that genetic deletion of *Spp1* improves disease severity in the orthologous *Pkd1*^RC/RC^ mouse model. We also created a freely available, searchable website (https://bmblx.bmi.osumc.edu/scPKD/) that can be used to identify cross- and intra-model specific changes in gene expression, guiding researchers to new therapeutic targets for treating PKD.

## Methods

### Single-cell RNA sequencing

This study combined previously published scRNAseq data^6–9^ with new data as outlined in supplemental methods. After sequencing, data was aligned to the reference genome (mm10) using Cellranger. The resulting matrix files were loaded into Seurat version 5.2.0 followed by removal of doublets (DoubletFinder^10^) and ambient RNA (Soupx^11^). Individual samples were merged using the ‘Merge’ function in Seurat followed by data integration using SCT-RPCA and Harmony^12^. Cells with over 50% of genes mapping to mitochondrial transcripts and unique gene counts greater than 3,000 or less than 200 were discarded, thresholds that have previously been established for kidney scRNAseq data^13^. After quality control measures and integration, we followed the standard Seurat vignette for clustering and data visualization. To identify differentially expressed genes in each cell cluster, we used the function ‘FindAllMarkers’ in Seurat on the normalized gene expression data. To find differences between control and PKD samples, we used the Wilcox rank sum text and ‘FindMarkers’ function in Seurat or DESeq2^14^ on data that was pseudobulked using the ‘AggregrateExpression’ function and the counts slot. Cell-cell communication was analyzed using CellChat^15^ while pathway and transcription factor inference was done using decoupleR^16^.

### Spatial transcriptomics

Spatial transcriptomics was performed on control and cystic samples from *Ift88* and *Pkd1*^RC/RC^ mice using 10X Visium as outlined in supplemental methods. Data was aligned to the reference genome using SpaceRanger followed by data processing using the standard vignette in scanpy. Spot deconvolution was done using the reference atlas and TACCO^17^ while cell-cell communication within a spatially confined distance was done using COMMOT^18^.

### Data and code availability

All raw scRNAseq and spatial transcriptomics data is available in GEO as outlined in supplemental methods. Processed and annotated .rds files for the whole atlas and each individual model can be downloaded from the website (https://bmblx.bmi.osumc.edu/scPKD/). Code used to generate figures is available in the lab’s github account (kzimmer1) under the repository name “Mouse-SingleCell-Atlas”.

## Results

### Generation of a cross-model, single-cell RNA sequencing atlas of mouse PKD

We analyzed scRNAseq data collected from six different mouse models of PKD, including published (Pax8 teto^cre^ *Pkd1*^f/f^ mice harvested 66, 100, or 130 days after doxycycline induction^7^; adult induced *Ift88* mice [induced at 8 weeks of age] harvested ∼7 months post induction^13^; adult induced *Pkd2* mice harvested ∼4 months post induction^9^; *Pkhd1*^cre^ *Pkd1*^f/f^ mice harvested at 7, 14 and 21 days of age^8^; and adult induced *Ift88* mice harvested 8 weeks post ischemia reperfusion injury^13^) and unpublished data (*Pkd1*^RC/RC^ mice harvested at one year of age; Figure 1A; harvest times depicted with a red arrow). A detailed description of grouping, genetic mutation, cell types impacted by the genetic mutation, time points of harvest, technology used, biological sex, total number of cells analyzed per sample, and PKD severity, when available, can be found in Supplemental table 1. After removal of doublets, ambient RNA, and low-quality cells (see supplemental methods), we clustered and annotated 265,644 cells from 41 samples (19 control; 22 PKD), identifying all major cell types including tubular epithelial cells, immune cells, microvasculature, fibroblasts, and podocytes (Figure 1B,C).

**Figure 1.**
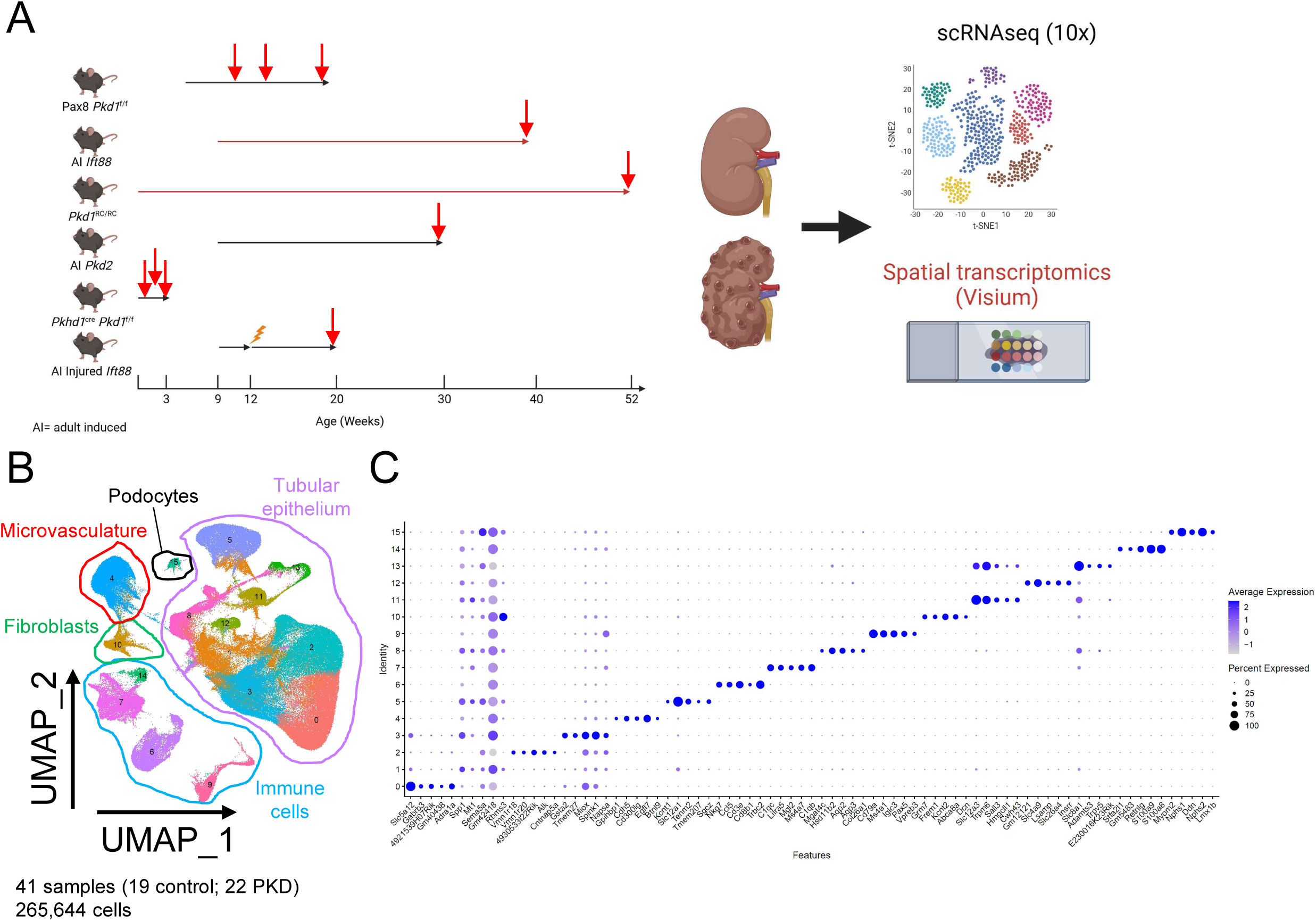
Single cell atlas of PKD in mice. **(A)** Schematic of experimental design used to generate the single cell atlas of PKD in mice. Red arrows indicate time points of harvest. **(B)** Uniform Manifold Approximation and Projection (UMAP) of all cells collected from mouse models of PKD. **(C)** Dotplot showing the top 5 markers of each cluster from the whole atlas. Pax8^cre^ *Pkd1*^f/f^ model (8 control samples, 9 PKD samples; all male), adult induced (AI) *Ift88* model (2 controls, 2 PKD; all female), *Pkd1*^RC/RC^ model (2 controls, 3 PKD; 3 males [1 control, 2 PKD], 2 females [1 control, 1PKD]), AI *Pkd2* (2 controls, 2 PKD; all female), *Pkhd1*^cre^ *Pkd1*^f/f^ (3 controls, 3 PKD; mix of males and females), AI injured *Ift88* (4 controls [2 sham operated *Ift88,* 2 injured cre negative control], 2 PKD; all female).

To better understand how mutations in PKD-related genes impacted tubular epithelium across models, we subsetted and re-clustered epithelial cells in isolation. Analysis of the data revealed 12 transcriptionally distinct clusters of cells spanning the entire length of the nephron including: proximal tubule (Clusters 0, 1, 2, 5), Loop of Henle (LOH; clusters 3, 7, 9), distal tubule (Clusters 6, 8, 11), and collecting duct (clusters 4, 10; Figure 2A,B). The proportion of S1-S3 proximal tubule, collecting duct principle cells, lower limb of the LOH, and thin descending limb of the LOH were increased in combined PKD samples compared to controls, with changes in the lower limb LOH being the most significant (Figure 2C,D). Across all nephron segments examined, tubular epithelial cells from PKD samples had increased expression of genes associated with tubular injury (*Lcn2, Havcr1*), complement (*C3*), inflammation (*Cxcl1*), and fibrosis (*Col1a1, Spp1, Sparc*), in agreement with previous literature (Figure 2E; Supplemental table 2)^19, 20^. Pathway and transcription factor inference revealed that several clusters expressed genes associated with pro-inflammatory pathways (JAK-STAT, NFκB, PI3K, TNF) and transcription factors (Figure 2F,G).

**Figure 2.**
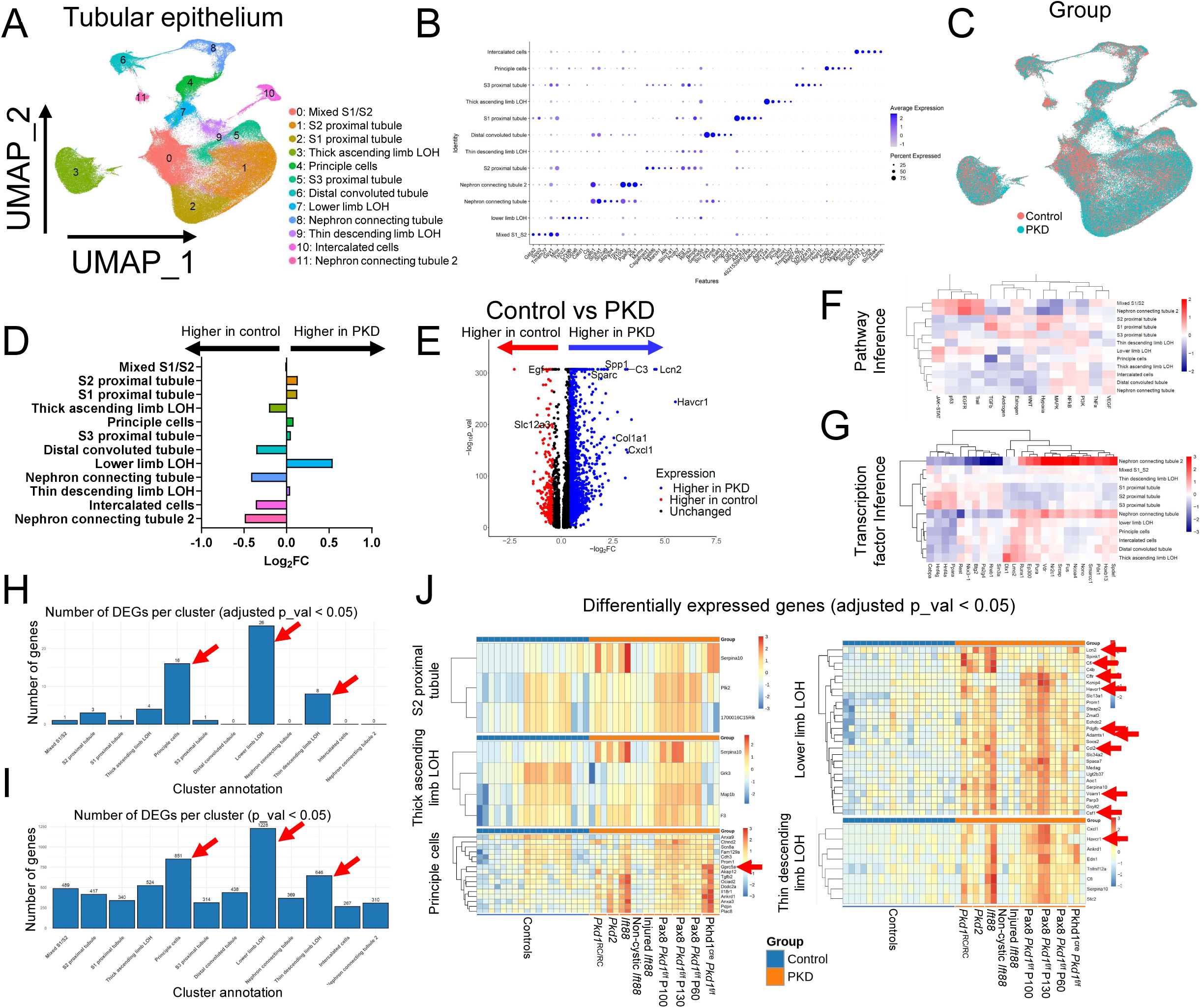
Analyses of scRNAseq data from the nephron. **(A)** UMAP of tubular epithelial cells subclustered from Figure 1. **(B)** Dotplot showing the top 5 markers of each cluster of cells. **(C)** UMAP of tubular epithelial cells based on experimental group, **(D)** Quantification of cluster abundance in control and PKD samples from whole atlas. **(E)** DEGs in all tubular epithelial cells when comparing control and PKD samples. **(F, G)** Pathway and transcription factor inference from DecoupleR. **(H,I)** Quantification of the number of DEGs in each cluster (control vs PKD) in pseudobulked scRNAseq data determined using DESeq2. **(J)** DEGS between control and PKD tubular cell clusters.

To better understand how PKD impacted gene expression in each nephron segment, we performed pseudobulk analysis of our scRNAseq data, followed by identification of differentially expressed genes (DEGs) in each cluster using DESeq2^14^. This analysis revealed that principle cells, lower limb LOH, and thin descending limb LOH clusters had the most DEGs between groups (Figure 2H,I), independent of the p value threshold that was used (adjusted p value < 0.05 or p value < 0.05). When we analyzed DEGs between control and PKD cells within each nephron segment (using adjusted p value < 0.05), we found that PKD samples had increased expression of genes associated with kidney injury (*Lcn2, Havcr1, Vcam1*), complement (*Cfi, C4b*), inflammation (*Csf1*, *Cxcl1, Ccl2*), fibrosis (*Pdgfb, Admats1*), and fluid/ion secretion (*Slc34a2, Slc13a1, Cftr, Kcnip4*; Figure 2J, red arrows). The majority of these DEGs were found in the lower limb LOH, although principle cells and cells from the thin limb LOH also expressed a significant number of genes previously associated with PKD. We also found that expression of *Gprc5a,* a recently identified marker of cyst lining epithelial cells^7^, was significantly increased in principle cells from PKD mice (Figure 2J).

When analyzing the data, we noted inconsistencies in gene expression across individual PKD replicates. To better understand this variability, we grouped DEGs (adjusted p value < 0.05) based on variables that may be driving this effect including sex, experimental model, time point of harvest, rate of disease progression, and genetic mutation (orthologous vs non-orthologous) and replotted the data using heatmaps. For rate of disease progression, mice were considered to have rapidly progressing PKD if they developed cysts in 8 weeks of fewer (*Pkhd1*^cre^ *Pkd1*^f/f^ model [3 samples]; Injured *Ift88* model [2 samples]). Independent of which nephron segment we analyzed, we found that DEGs segregated based on the rate of disease progression rather than sex, experimental model, time point of harvest, or genetic mutation (Supplemental figures 1-3), suggesting that rate of PKD progression, and not the type of genetic mutation, most heavily impacts gene expression.

**Figure 3.**
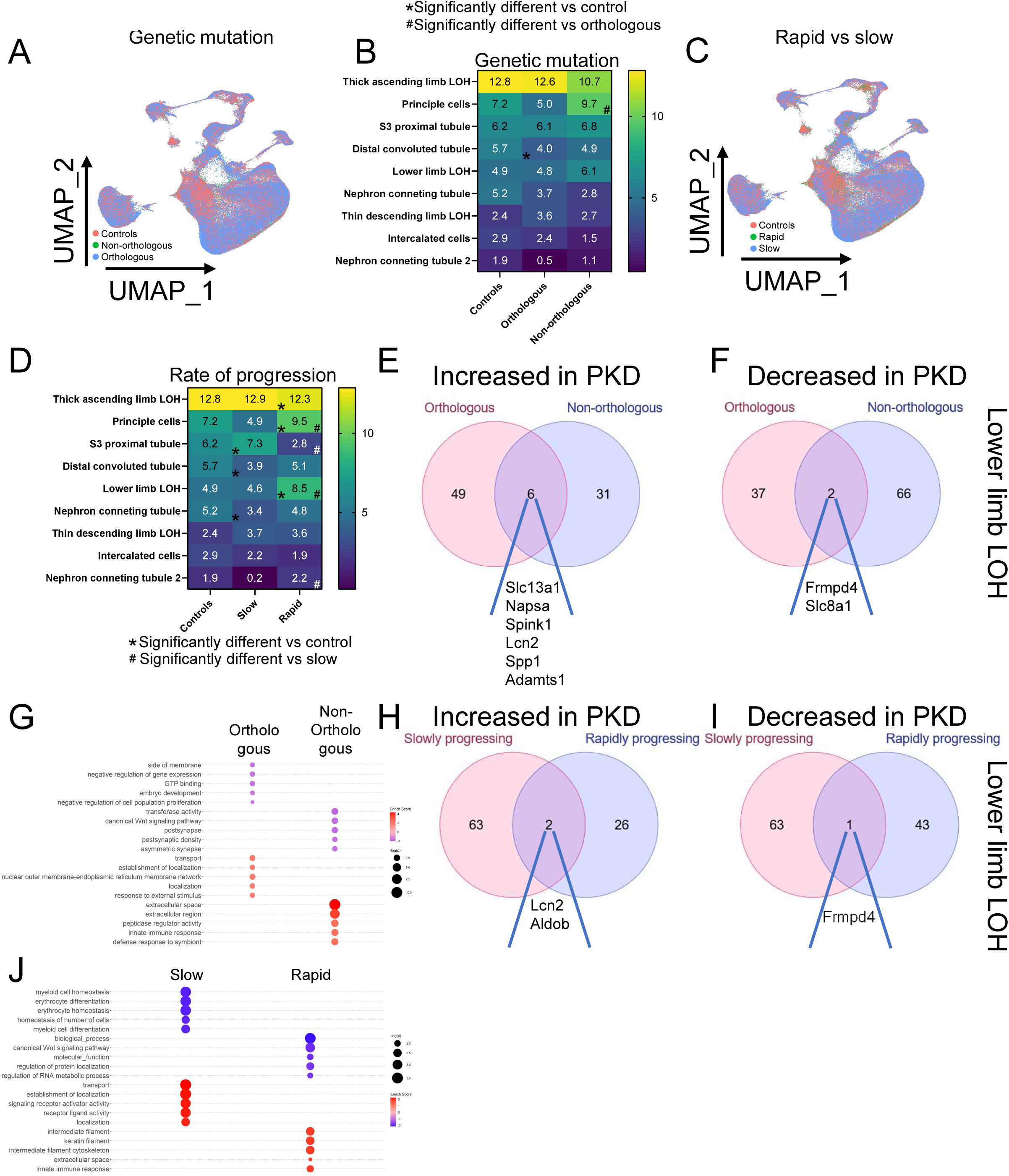
Analysis of cluster abundance and gene expression based on genetic mutation and rate of PKD progression. **(A)** UMAP of tubular epithelial cells based on genetic mutation (orthologous vs non-orthologous mutation). **(B)** Quantification of cluster abundance based on genetic mutation. **(C)** UMAP of tubular epithelial cells based on rate of PKD progression (slow vs rapid). **(D)** Quantification of cluster abundance based on rate of PKD progression. **(E,F)** Venn diagram showing unique and shared DEGs that are **(E)** increased or **(F)** decreased in the lower limb LOH based on the genetic mutation. **(G)** Bubble plot of the top 5 GO pathways that are increased or decreased in lower limb LOH based on genetic mutation. **(H,I)** Venn diagram showing unique and shared DEGs that are **(H)** increased or **(I)** decreased in the lower limb LOH based on the rate of PKD progression. **(J)** Bubble plot of the top 5 GO pathways that are increased or decreased in lower limb LOH based on the rate of PKD progression.

### The impact of the type of genetic mutation and the rate of PKD progression on cluster abundance and gene expression

To understand how the genetic mutation impacts cluster abundance in our atlas, we created UMAP projections based on the genetic mutation and re-quantified cluster abundance (Figure 3A). For this quantification, we excluded samples from the S1 and S2 proximal tubule as we found that the sequencing technology (single cell vs single nucleus) heavily impacted cluster abundance, independent of phenotype (Supplemental figure 4). Quite surprisingly, analysis of cell proportions based on the genetic mutation (orthologous vs non-orthologous) showed minimal differences between groups, with only principle cells being significantly increased in non-orthologous models compared to orthologous models and distal convoluted tubule being decreased in orthologous models compared to non-cystic controls (Figure 3A,B). We also analyzed how cluster proportions were altered based on the rate of PKD progression as this was the metadata variable that most heavily influenced DEGs when comparing control to PKD samples (Supplemental figures 1-3). Quantification of cluster proportions revealed significant differences in several cell types including the thick ascending limb LOH, principle cells, S3 proximal tubule, distal convoluted tubule, lower limb LOH, nephron connecting tubule, and nephron connecting tubule 2 (Figure 3C,D).

Next, we set out to understand how the genetic mutation and rate of PKD progression impacted gene expression across nephron segments. To do this, we pseudobulked single cell data from each nephron segment based on the appropriate metadata variable followed by performing DESeq2 on pairwise data (control vs orthologous, control vs non-orthologous, etc). We then compared genes that were differentially expressed in each group vs controls using a Venn diagram, with overlapped genes representing ones that were increased or decreased in both groups vs non-cystic controls. When we did the analysis based on the genetic mutation, we found that six out of a possible 37 genes (16%) were upregulated (*Slc13a1, Napsa, Spink1, Lcn2, Spp1, Adamts1*) and that two out of a possible 39 genes (5%) were downregulated (*Frmpd4, Slc8a1*) in both groups vs non-cystic controls (Figure 3E,F; Supplemental figure 5A,B, Supplemental table 3). We confirmed that expression of shared genes were increased/decreased in both groups compared to non-cystic controls at the single cell level (Supplemental figure 5C,D). Of note, when we analyzed expression of shared genes at the individual model level, we found that changes in gene expression were often driven by one or two mouse models, the most frequent being the *Ift88* and *Pkd1*^RC/RC^ models (Supplemental figure 6E,F). Pathway analysis of genes enriched in either group alone compared to non-cystic controls revealed that orthologous models were associated with transport, localization, and response to stimuli while non-orthologous models were associated with extracellular space and innate immune response (Figure 3G).

When we analyzed DEGs in the lower limb LOH based on the rate of disease progression, we found that only two out of a possible 28 genes (7%) were increased (*Lcn2, Aldob*) and that one out of a possible 44 genes (2%) were decreased (*Frmpd4*) in both groups compared to non-cystic controls (Figure 3H,I; Supplemental figure 6A,B; Supplemental table 4). Once again, we confirmed that the selected genes were increased/decreased in both groups at the single cell level (Supplemental figure 6C,D) and that changes in gene expression were often driven by one or two mouse models (Supplemental figure 6E,F). Pathway analysis of genes enriched in either group alone compared to non-cystic controls showed that slow PKD models had increased expression of genes associated with transport, localization, and receptor-ligand activity while rapid models had increased expression of genes associated with filaments and cytoskeleton (Figure 3J). Thus, we concluded that the genetic mutation has less impact on gene expression than the rate of PKD progression.

### Model specific differences in gene expression across nephron segments

Throughout our analyses, we consistently noted that DEGs were driven by one or two individual models. As such, we analyzed how cluster proportion and DEGs were different in PKD versus control samples from each individual model (Figure 4A). Once again, we excluded S1-S2 proximal tubule segments from our analyses due to technology specific differences that were observed. When we analyzed the data, we found differences in cluster proportion in principle cells, S3 proximal tubule, distal convoluted tubule, lower limb LOH and nephron connecting tubule when comparing control to PKD samples within individual models, with the injured *Ift88*, *Pkd2*, and *Pkd1*^RC/RC^ models having the most differences in cluster proportions (Figure 4B). Cluster proportion was also most frequently and consistently affected in distal convoluted tubule (decreased in PKD vs controls) and lower limb LOH (increased in PKD vs controls) when comparing control to PKD samples (Figure 4B).

**Figure 4.**
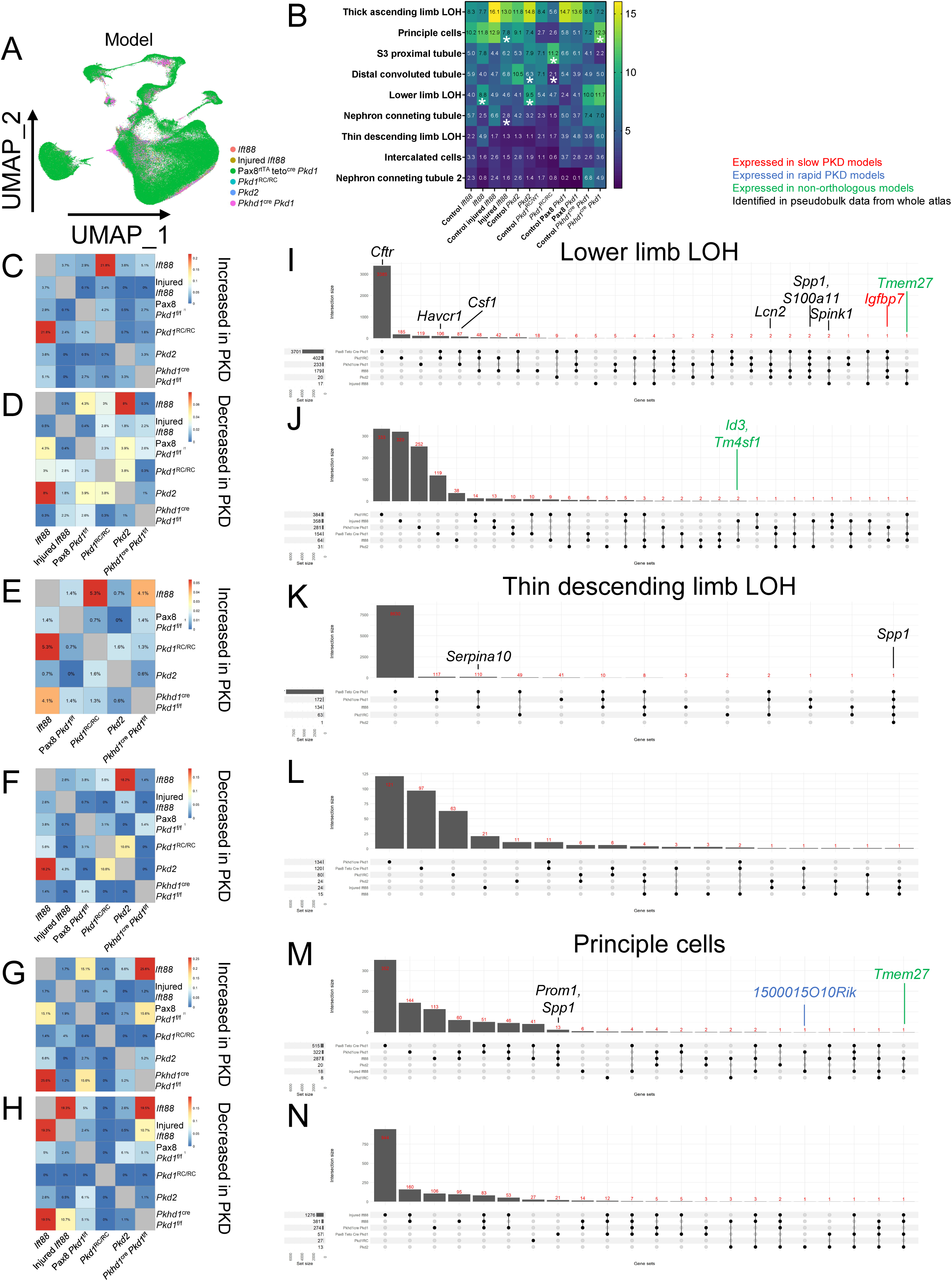
Model level analysis of cluster abundance and gene expression reveals consistent upregulation of *Spp1* in PKD. **(A)** UMAP of tubular epithelial cells based on individual models. **(B)** Quantification of cluster abundance in individual models. **(C-H)** Jaccard indices of intersected genes that were increased or decreased in the lower limb LOH, thin descending limb LOH, and principle cells of individual PKD models. **(I-N)** Upset plot showing shared genes that are increased or decreased across models. Genes highlighted in red are increased in all slow PKD models; genes highlighted in blue are increased in all rapid PKD models; genes highlighted in green are enriched in all non-orthologous models; genes highlighted in black were identified in pseudobulk analysis.

We next calculated DEGs (adj_p_val < 0.05) in each cystic model in relation to non-cystic controls from the same model using the Wilcox Rank Sum test and plotted the data using upset plots and Jaccard Indices to quantify model similarity. We focused our analyses on the lower limb LOH, thin descending limb LOH and principle cells due to the fact that these clusters had the highest number of DEGs at the whole atlas level. Quite surprisingly, we found that DEG overlap across all models was relatively limited, ranging from no overlap to 25.6% of DEGs, despite the fact that all models contained cysts at the time of harvest (Figure 4C-H). We also found that model similarity was different depending on the nephron segment being analyzed (Figure 4C-H). For example, hen comparing DEG overlap in the lower limb LOH, we found that the *Ift88* and *Pkd1*^RC/RC^ models had the most overlap in increased DEGs (i.e. genes were increased compared to controls in both models) while *Ift88* and *Pkd2* mice were most similar in terms of decreased DEGs (Figure 4C,D). Thus, while there are similarities across models, each model is unique in terms of DEGs that are altered across nephron segments.

At the individual model level, we found minimal overlap in DEGs across metadata variables (rate of progression, orthologous vs non-orthologous), independent of whether the gene was increased or decreased. For example, in the lower limb LOH, only *Igfbp7* was increased in all slow PKD models relative to non-cystic controls while only *Tmem27* was increased in all non-orthologous models relative to non-cystic controls (Figure 4I; Supplemental figure 7). Similarly, we found limited overlap of DEGs in cystic mice relative to non-cystic controls across all models, with the exception of *Spp1,* which was increased in five out of six PKD models in the lower limb LOH and thin descending limb LOH (Figure 4I,K). While several other DEGs identified in pseudobulk analysis of the whole atlas failed to be increased in all models relative to non-cystic controls, several of the DEGs were conserved in multiple models (Figure 4I-N). For example, *Lcn2*, which was enriched in PKD samples in the lower limb LOH in the whole atlas (Figure 2J), was increased in four out of six models relative to non-cystic controls (Figure 4I; Supplemental figure 7E). We also noted that the number of increased and decreased DEGs was highly variable between models and nephron segments (Figure 4I-N). A breakdown of model specific genes that were increased or decreased in other nephron segments can be found in Supplemental figures 8-10. In summary, we find that there are limited DEGs that are shared across models of PKD, likely due to subtle differences in model specific controls (age of harvest, etc), with the exception of *Spp1,* which was highly enriched in multiple PKD nephron segments.

### Identification of PKD specific clusters in the lower limb LOH and thin descending limb LOH

When analyzing cluster abundance and DEGs, we initially grouped all cells from each nephron segment into one homogenous population, without making assumptions about PKD specific subsets that may be embedded within these nephron segments. To identify possible PKD-specific clusters, we subsetted and re-clustered each nephron segment at high resolution, looking for subclusters that were highly enriched in PKD samples in relation to non-cystic controls. This analysis revealed that two nephron segments, the lower limb LOH and thin descending limb LOH, each contained a single subsetted cluster that was over four-fold enriched in PKD samples compared to controls (Figure 5A). As such, we subsetted these two nephron segments for further analysis. When we did this, we found that two subsetted clusters in each nephron segment were greater than two-fold enriched in PKD samples relative to non-cystic controls (Figure 5B-E). These segments, which we annotated as lower limb PKD cluster 1, lower limb PKD cluster 2, thin limb PKD cluster 1, and thin limb PKD cluster 2, had highly disparate enrichment amongst individual PKD models (Figure 5F-I). For example, lower limb PKD cluster 1 was almost exclusively derived from the Pax8 *Pkd1*^f/f^ model while lower limb PKD cluster 2 was derived from the *Ift88, Pkd1*^RC/RC^, and *Pkhd1*^cre^ *Pkd1*^f/f^ models (Figure 5H). In contrast, the PKD enriched clusters in the thin descending limb LOH were derived from multiple models, although the frequency of each PKD cluster varied between models (Figure 5I). DEGs and pathways in each of the PKD enriched clusters from the lower limb were unique, with lower limb PKD cluster 1 expressing *Cftr* and having an enrichment of genes associated with the membrane and organonitrogen compound processes while lower limb PKD cluster 2 expressed *Spp1* and genes associated with development, differentiation, and signaling (Figure 5J,K). Thin limb PKD cluster 1 had enriched expression of *Spp1* and *Cftr*, which was associated with developmental processes while thin limb PKD cluster 2 had enriched expression of *Slc5a2,* which was associated with the membrane and amino acid transport (Figure 5L,M). Of note, *Spp1* was only enriched in the lower limb PKD cluster 2 and thin limb PKD cluster 1.

**Figure 5.**
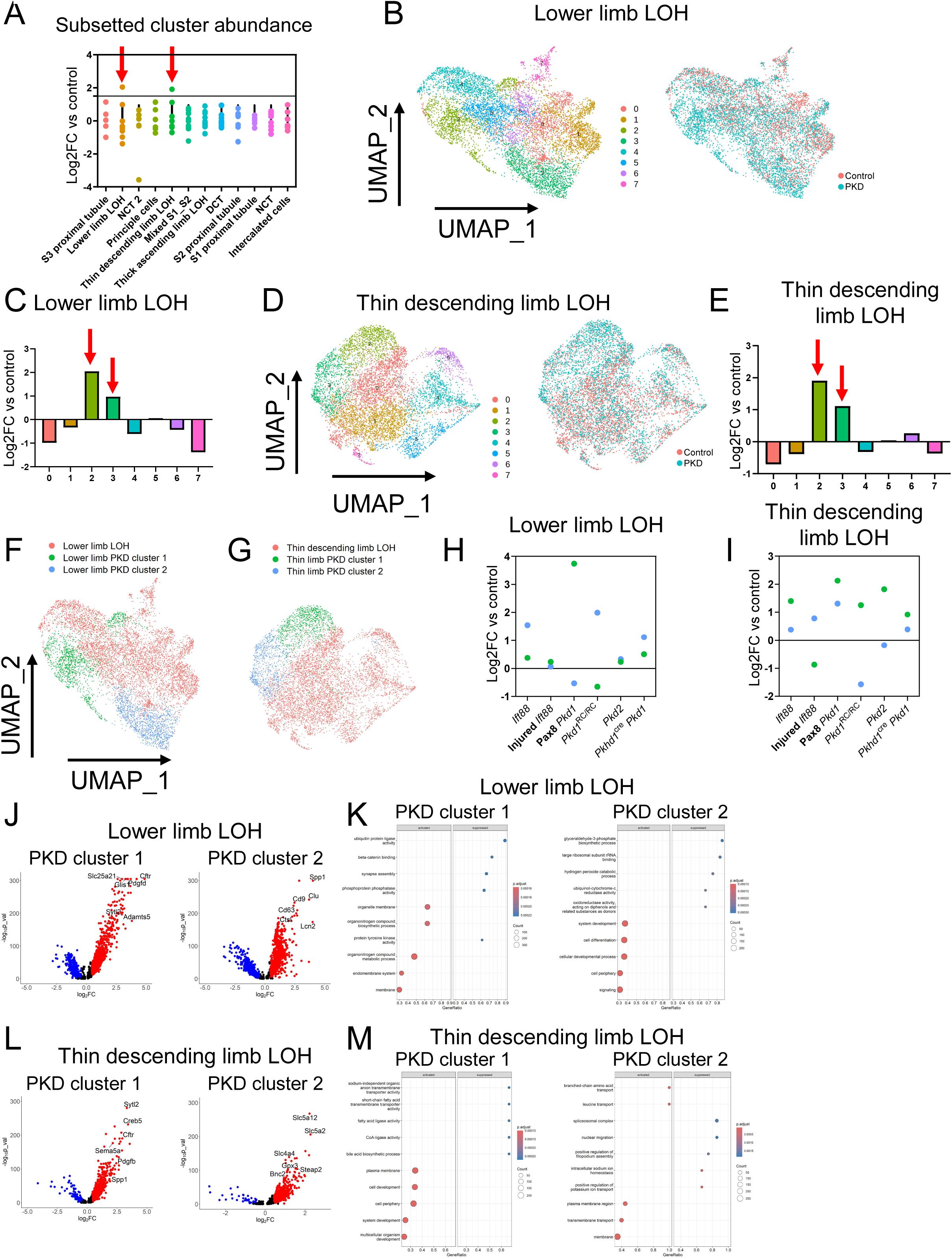
PKD enriched clusters across mouse models. **(A)** Quantification of subsetted, high resolution cluster abundance in each nephron segment. Data is plotted as Log2FC in relation to non-cystic controls. **(B)** UMAP showing high resolution clusters and groups in lower limb LOH. **(C)** Quantification of subsetted, high resolution lower limb LOH cluster. Two segments were greater than two-fold increased in PKD samples. **(D)** UMAP showing high resolution clusters and groups in thin descending limb LOH. **(E)** Quantification of subsetted, high resolution thin descending limb LOH cluster. Two segments were greater than two-fold increased in PKD samples. **(F,G)** Re-annotated high resolution clusters from **(F)** lower limb LOH and **(G)** thin limb LOH. **(H,I)** Quantification of PKD-enriched cluster abundance in **(H)** lower limb LOH and **(I)** thin limb LOH from individual models in relation to model specific, non-cystic controls. **(J,K)** Volcano plot and gene set enrichment analysis (GSEA) of genes expressed in each PKD- specific cluster from the lower limb LOH. **(L,M)** Volcano plot and gene set enrichment analysis (GSEA) of genes expressed in each PKD-specific cluster from the thin limb LOH.

### Atlas and model level analysis of cell-cell communication show consistent enrichment of SPP1 signaling

Our data indicate that the lower limb LOH, thin descending limb LOH, and principle cells are the nephron segments most altered in PKD. Further, by subsetting and re-clustering at high resolution, we were able to identify two PKD specific clusters in both the lower limb LOH and thin limb LOH. To understand whole atlas and model level signaling between PKD-specific clusters and other cell types, we performed CellChat on the fully annotated single cell atlas (Supplemental figure 11). In the whole atlas, we found that the number and strength of interactions was increased between lower limb PKD cluster 1 and fibroblasts in PKD samples (Supplemental figure 12A). The strength, albeit not the number, of interactions was also increased in PKD samples from lower limb PKD cluster 2 and thin limb PKD cluster 1 in the whole atlas (Supplemental figure 12A). When we analyzed signaling interactions at the model level, we found that each model was unique in regards to the cluster with the greatest number and strength of interactions (Supplemental figure 12B-G). For example, we found that thin limb PKD cluster 2 had the most frequent and strongest interactions in *Ift88,* injured *Ift88, Pkd1*^RC/RC^, and *Pkhd1*^cre^ *Pkd1*^f/f^ mice, although predicted interaction partners were different in each model (Supplemental figure 12B,C,E,G). We also found that lower limb PKD cluster 1 had the most frequent and strongest interactions in the Pax8 *Pkd1*^f/f^ and *Pkd2* models, although signaling partners were once again distinct between models (Supplemental figure 12D,F).

We next analyzed incoming and outgoing signaling in PKD enriched clusters at the atlas and model level. Analysis of the data revealed that SPP1 was the strongest outgoing and incoming signaling pattern in the whole atlas and in each individual model (red arrow), with the exception of incoming signaling in the *Pkhd1*^cre^ *Pkd1*^f/f^ model (Figure 6A-N). With regards to cell types, we found that lower limb PKD cluster 2 had the highest outgoing signaling strength across models while the cell type receiving the strongest signal varied (Figure 6A-N). Other outgoing signaling pathways of interest included COMPLEMENT, CSF, SEMA3, WNT, and PDGF, several of which have been associated with PKD in the past^21–24^. Other incoming signaling pathways of note included GALECTIN, IGF, TWEAK, EGF, and MIF, which have also been associated with PKD ^25–29^.

**Figure 6.**
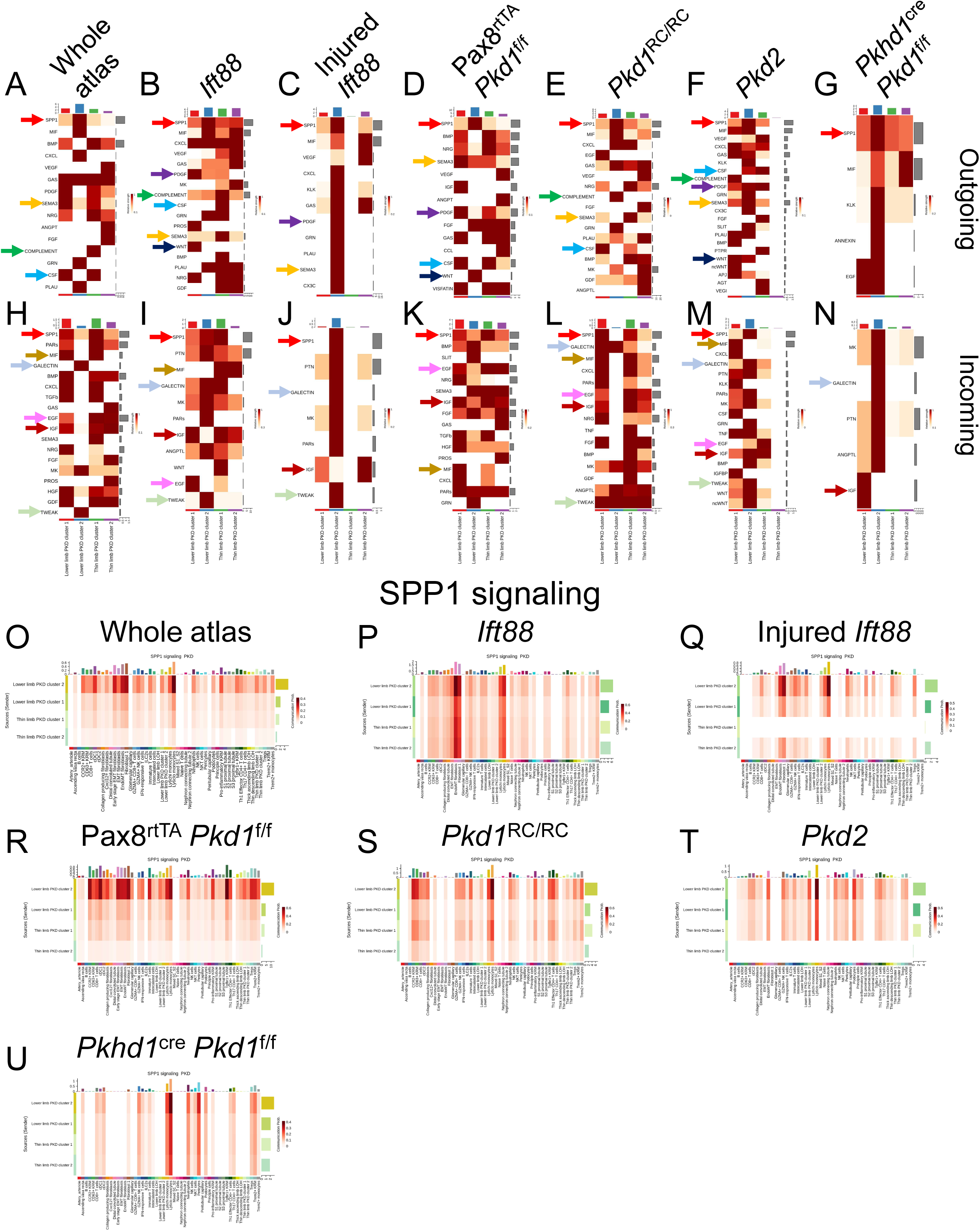
Cellchat analysis of cell-cell communication in PKD-enriched clusters at the whole atlas and model level reveals elevated SPP1 signaling to monocytes. (A-G) Top outgoing signaling pathways from PKD enriched clusters at the whole atlas and model level. Arrows indicate pathways that are conserved across multiple models. **(H-N)** Top incoming signaling pathways to PKD enriched clusters at the whole atlas and model level. Arrows indicate pathways that are conserved across multiple models. **(O-U)** SPP1 signaling from PKD enriched clusters to all other cell types at the whole atlas or model specific level.

Throughout our analyses, we consistently found that SPP1 signaling, both incoming and outgoing, was increased in the PKD-enriched clusters from the lower limb LOH and thin limb LOH. A more thorough analysis of outgoing SPP1 signaling from those cell types revealed frequent communication with Ly6c^hi^ and Ly6c^lo^ monocytes, in line with data indicating that SPP1 serves as a monocyte chemoattractant (Figure 6O-U)^30, 31^. We also noted that outgoing SPP1 signaling from *Ift88,* injured *Ift88,* and Pax8 *Pkd1*^f/f^ mice was predicted to interact with fibroblasts in these models. Thus, while increased SPP1 expression and signaling is a clear hallmark of PKD, our data suggest that signaling partners may be model specific.

### Spatial transcriptomics shows that SPP1 signaling is highly enriched in *Pkd1*^RC/RC^, but not *Ift88,* kidneys

To understand if dysregulated signaling networks found in PKD samples using scRNAseq are present within spatially restricted niches, we subjected control and cystic *Pkd1*^RC/RC^ and *Ift88* mice (2 control, 2 PKD from each model) to spatial transcriptomics using Visium 10X Genomics (Figure 7A; Supplemental figure 13, Supplemental methods). To achieve cell-type resolution, we used our fully annotated scRNAseq atlas (Supplemental figure 11) to deconvolute spots using TACCO^17^ and found that segments mapped to appropriate anatomical locations in control and PKD kidneys (Figure 7B,C; Supplemental figures 14-21). To test if the PKD-enriched clusters found in scRNAseq data were cyst localized, we outlined cysts using ImageJ and quantified the frequency with which each PKD-enriched cluster was found in cystic or non-cystic regions (Figure 7D,E). The data indicate that the proportion of lower limb PKD cluster 1 and thin limb PKD cluster 1 were enriched in cystic vs non-cystic regions, while lower limb PKD cluster 2 and thin limb PKD cluster 2 were equally dispersed between cystic and non-cystic regions (Figure 7F). Next, we analyzed cell-cell communication within a spatially restricted distance (200μM) using COMMOT^18^ and plotted the level of signaling interaction in PKD samples in relation to sex-matched, non-cystic controls from each model. We found that several of the signaling pathways found in our Cellchat analysis of scRNAseq data from *Ift88* and *Pkd1*^RC/RC^ mice were also enriched in spatial transcriptomics data (Figure 7G,H). For example, we found that WNT, COMPLEMENT, and CSF signaling were increased in all PKD enriched clusters of *Ift88* mice (Figure 7G) while *Pkd1*^RC/RC^ mice had enriched SPP1 and COMPLEMENT signaling in both scRNAseq and spatial transcriptomics data (Figure 7H). In contrast to scRNAseq CellChat data, we did not find enriched SPP1 signaling in *Ift88* mice when analyzing spatial transcriptomics data, likely due to the variability in *Spp1* expression between the two *Ift88* samples (Figure 7G; Supplemental figure 22).

**Figure 7.**
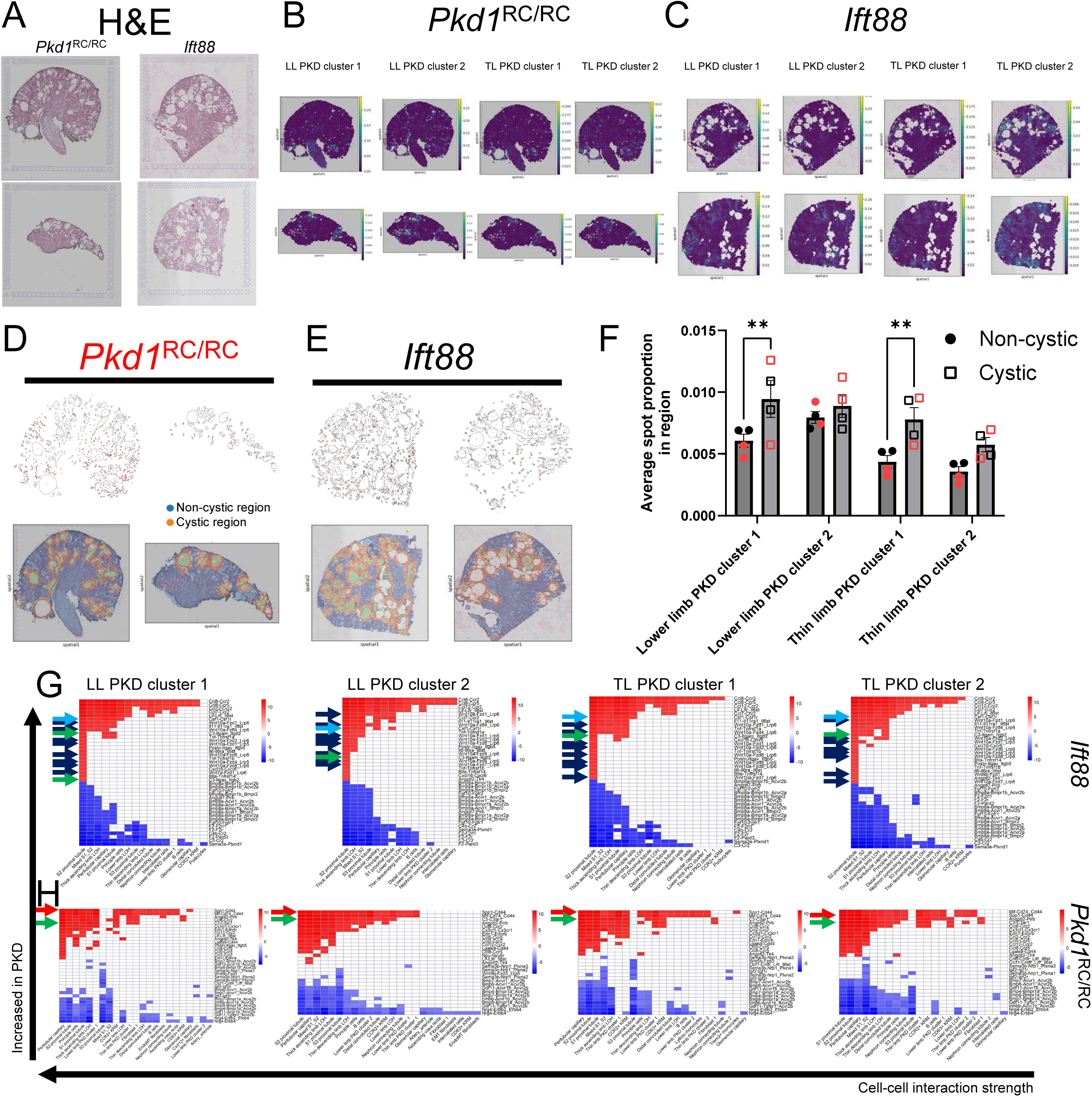
Spatial transcriptomics shows that SPP1 signaling is increased in the *Pkd1*^RC/RC^, but not *Ift88* model. **(A)** H&E images of PKD samples that were subjected to spatial transcriptomics. **(B,C)** TACCO deconvolution of PKD enriched clusters in **(B)** *Pkd1*^RC/RC^ and **(C)** *Ift88* mice. **(D,E)** Cystic and non-cystic regions in *Pkd1*^RC/RC^ and *Ift88* kidneys identified using ImageJ. **(F)** Quantification of the proportion of each PKD-enriched cluster that was found in cystic or non-cystic regions in *Pkd1*^RC/RC^ and *Ift88* mice. Proportions from *Pkd1*^RC/RC^ mice are shown in red. **(G,H)** Top 50 ligand-receptor pairs between PKD enriched clusters and other cell types. Data was normalized to non-cystic kidneys.

We also investigated cell-cell signaling in cystic regions by selected all spots that directly touched outlined cysts plus one spot on the basolateral side (Supplemental figure 23A). Quite surprisingly, when we did this analysis, we found that a majority of signaling interactions that were enriched in PKD samples when analyzing whole kidney spatial data or scRNAseq data were not increased in cystic regions in either model, with the exception of SEMA3 signaling (Supplemental figure 23B,C). SPP1 signaling was also not increased in cystic regions of *Pkd1*^RC/RC^ mice although it was increased when comparing PKD to control kidneys, suggesting that SPP1 dysregulation in PKD is not cyst specific.

### Cyst size does not impact highly variable or differentially expressed genes

It is commonly believed that the phenotype, and likely the transcriptional signature, of cystic epithelium is different depending on the size of the cyst. To test this idea, we quantified cyst size across all four spatial transcriptomics PKD replicates, binned them into small, medium, or large sized cysts, and analyzed the top 100 highly variable genes (HVGs) across all spots (Supplemental figure 24A). Surprisingly, in contrast to our initial hypothesis, we found that cyst size did not impact HVGs in either the *Pkd1*^RC/RC^ or *Ift88* model (Supplemental figure 24B). Likewise, when we analyzed DEGs between different sized cysts within experimental models, we found minimal differences (Supplemental figure 24C), although we were able to identify some differences when comparing different sized cysts to normal spots (Supplemental figure 25). Thus, we concluded that cyst size does not heavily influence gene expression, at least in our spatial transcriptomics data.

### Individual spots in the *Ift88* and *Pkd1*^RC/RC^ kidneys have an injury, inflammation, and fibrosis signature

To better understand factors that may driving the previously reported heterogeneity observed in PKD kidneys, we isolated individual cysts using ImageJ (Supplemental figure 26A). When we analyzed HVGs, we found significant heterogeneity at both the cyst and spot level (Supplemental figure 26B,C). While *Pkd1*^RC/RC^ kidneys had individual cysts that expressed genes associated with injury (*Lcn2, Havcr1*), inflammation (*C1qa, Ccr2, Spp1*), and fibrosis (*Fn1, Col1a1, Col3a1*), we were unable to find cysts with this signature in the *Ift88* model (Supplemental figure 26B,C). However, both the *Pkd1*^RC/RC^ and *Ift88* model had individual spots expressing genes associated with injury, inflammation, and fibrosis (Supplemental figure 26C), although the signature was more pronounced in the *Pkd1*^RC/RC^ model. When we mapped cysts and spots with an injury, inflammation, and fibrosis signature back to the spatial data, we found several cysts in the *Pkd1*^RC/RC^ model that were entirely encompassed by the injury, inflammation, and fibrosis signature whereas this signature was diffusely scattered throughout cystic regions in the *Ift88* model (Supplemental figure 27).

### Loss of *Spp1* improves PKD in the orthologous *Pkd1*^RC/RC^ model

When analyzing scRNAseq and spatial transcriptomics data, we found that SPP1 expression and signaling was consistently enriched in the *Pkd1*^RC/RC^ model of PKD. In contrast, *Ift88* mice showed enriched SPP1 expression and signaling, but only in scRNAseq data. Thus, we further investigated SPP1 interactions in the *Pkd1*^RC/RC^ model. When we did this, we found that *Spp1* expression was strongly increased in multiple nephron segments in *Pkd1*^RC/RC^ mice including lower limb PKD cluster 2 and thin limb PKD cluster 1 (Figure 8A), in agreement with Cellchat data showing that those clusters had the strongest outgoing SPP1 signaling in *Pkd1*^RC/RC^ mice. *Spp1* expression was also increased in *Pkd1*^RC/RC^ kidneys compared to control kidneys as determined by spatial transcriptomics and immunohistochemistry (Figure 8B-D). Of note, *Spp1* (OPN) was expressed in both cystic and non-cystic regions in *Pkd1*^RC/RC^ mice (Figure 8D, Supplemental figure 28), once again indicating that *Spp1* expression is increased throughout PKD kidneys and is not centralized to cystic regions. When analyzing ligand-receptor pairs in scRNAseq and spatial transcriptomics data, we consistently observed increased *Spp1-Itga4/Itgb1* signaling between lower limb PKD cluster 2, thin limb PKD cluster 1, and Ly6c^lo^ monocytes (Supplemental figures 29,30).

**Figure 8.**
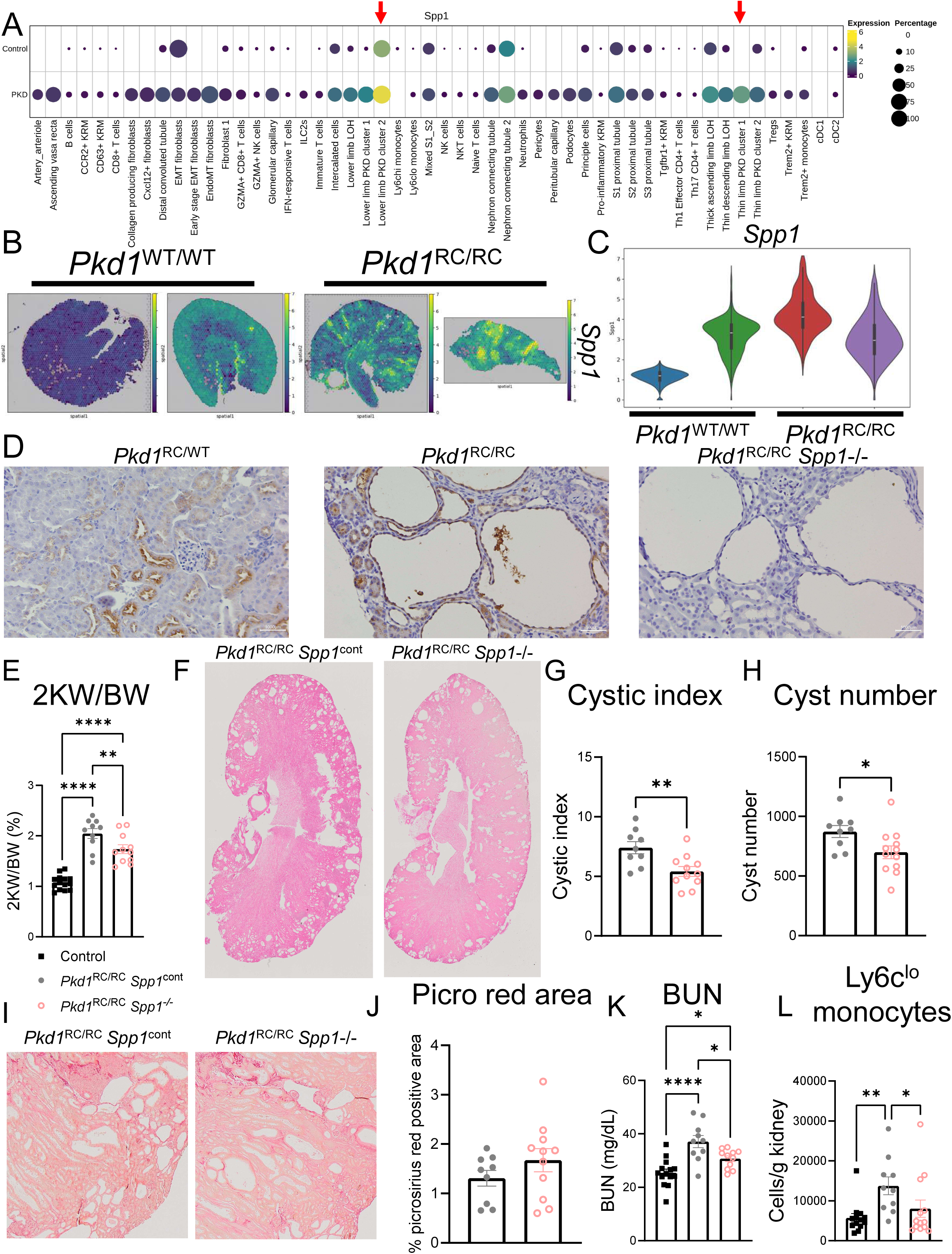
Loss of *Spp1* improves PKD severity and reduces Ly6c^lo^ monocyte number in the *Pkd1*^RC/RC^ model. **(A)** Dotplot showing expression of *Spp1* in all cell types of *Pkd1*^RC/RC^ mice. **(B)** *Spp1* expression in *Pkd1*^WT/WT^ and *Pkd1*^RC/RC^ sections. **(C)** Quantification of *Spp1* expression in *Pkd1*^WT/WT^ and *Pkd1*^RC/RC^ sections. **(D)** Immunohistochemistry staining of osteopontin in *Pkd1*^RC/WT^ and *Pkd1*^RC/RC^ sections. Images were taken using 20X magnification. **(E)** Quantification of two kidney weight to body weight (2KW/BW) ratio in *Pkd1*^RC/RC^ *Spp1*^cont^ and *Pkd1*^RC/RC^ *Spp1*-/- mice at one year of age. **(F-H)** H&E stained images of *Pkd1*^RC/RC^ *Spp1*^cont^ and *Pkd1*^RC/RC^ *Spp1*-/- mice along with quantification of **(G)** cystic index and **(H)** cyst number. **(I,J)** Picrosirius red stained sections **(I)** and quantification **(J)**. **(K)** Quantification of kidney function as measured by blood urea nitrogen (BUN). **(L)** Flow cytometry quantification of Ly6c^lo^ monocyte number. One way ANOVA (B,I), Student’s T-test (all others). *P <0.05, **P <0.01; ***P <0.001; ****P <0.0001. Control (*Pkd1*^RC/WT^) *Spp1*^cont^= 14 mice, *Pkd1*^RC/RC^ *Spp1*^cont^= 10 mice, *Pkd1*^RC/RC^ *Spp1*-/-= 12 mice; 36 total *Pkd1*^RC/RC^ mice.

To test the functional importance of *Spp1* in PKD, we crossed *Pkd1*^RC/RC^ mice to the *Spp1* knockout mice and analyzed PKD severity at ∼1 year of age. Analyses of one-year old *Pkd1*^RC/RC^ mice showed that loss of *Spp1* improved 2KW/BW, cystic index, and cyst number compared to aged matched, littermate controls (Figure 8E-H). Despite the fact that no quantifiable differences were found in fibrosis between groups as indicated by picrosirius red staining, kidney function was improved in *Pkd1*^RC/RC^ *Spp1* knockout mice compared to *Pkd1*^RC/RC^ *Spp1* control mice (Figure 8I-K). Loss of *Spp1* also reduced the number of Ly6c^lo^ monocytes in *Pkd1*^RC/RC^ mice (Figure 8L), while other immune cell numbers remained unchanged (Supplemental figure 31).

## Discussion

In this manuscript, we generated a comprehensive scRNAseq atlas comprised of 6 different mouse models of PKD whereby we provide a detailed description of genes, pathways, and signaling interactions that are disrupted across and within individual PKD models and cell types. Second, we performed spatial transcriptomics in two mouse models of PKD and found that several of the dysregulated signaling pathways identified via scRNAseq, were also present ins spatial transcriptomics data. Quite surprisingly, despite its consistent enrichment in scRNAseq data across and within individual models, spatial transcriptomics data indicate that SPP1 expression and signaling was only enriched in the *Pkd1*^RC/RC^, but not *Ift88* model of PKD. Using the integrated atlases, we found that SPP1 was predicted to interact with Ly6c^lo^ monocytes through the *Itga4-Itgb1* receptor complex. Loss of *Spp1* resulted in improved PKD severity and reduced Ly6c^lo^ monocyte number, suggesting that SPP1 promotes PKD progression by promoting Ly6c^lo^ monocyte recruitment in an *Itga4-Itgb1* dependent manner. To facilitate the use of these data amongst the PKD community, we developed a user-friendly and freely accessible website (https://bmblx.bmi.osumc.edu/scPKD/) whereby researchers can query genes of interest in the whole scRNAseq atlas or in individual mouse models. We have also provided a visual step-by-step guide for navigating the website in Supplemental figure 32.

The epistatic relationship between primary cilia and polycystin proteins is well-established. Inherent to this concept is the idea that gene expression is unique when comparing mice lacking primary cilia or polycystin proteins^32^. Quite surprisingly, when we analyzed our cross-model atlas, we found that DEGs did not segregate based on the type of genetic mutation (orthologous vs non-orthologous), but instead segregated based on the rate of PKD progression. In line with these data, we found that the rate of PKD progression had a greater impact on cluster abundance and gene expression when compared to the type of genetic mutation.

Another interesting observation from our manuscript was the predicted interaction between PKD-enriched clusters and Ly6c^lo^ monocytes through a *Spp1*/*Itga4-Itgb1* signaling axis. In support of this predicted interaction, we found that loss of *Spp1* in the *Pkd1*^RC/RC^ model significantly reduced the number of Ly6c^lo^ monocytes in the kidney, while all other immune cell subsets analyzed were unchanged. The fact that our computational approaches predicted an interaction between PKD-enriched clusters and Ly6c^lo^ monocytes through *Spp1*/*Itga4-Itgb1* signaling, combined with paralleling data showing reduced PKD severity and Ly6c^lo^ monocyte numbers, suggest that Spp1 promotes PKD progression through *Itga4-Itgb1* dependent, Ly6c^lo^ monocyte recruitment and/or function. This is further supported by published data showing that loss of *Itgb1* reduced PKD severity in an orthologous *Pkd1* model of disease^33^. Further investigation of Ly6c^lo^ monocyte involvement in PKD is warranted.

In conclusion, we generated an integrated single-cell and spatial atlas of cystic kidney disease across multiple PKD mouse models. Using this atlas, we identified dysregulated SPP1 signaling that was conserved across mouse models and show that loss of *Spp1* improved PKD severity in the orthologous *Pkd1*^RC/RC^ model, possibly through *Itga4-Itgb1* dependent Ly6c^lo^ monocyte recruitment.

## Disclosure statement

The authors have nothing to disclose.

## Acknowledgments

We thank the Institutional Research Core Facility at OUHSC for the use of the Core Facility which provided imaging, spatial library generation, and sequencing.

## Funding

These studies were supported in part by the following research grants: Polycystic Kidney Disease Research Foundation grant 826369 (KAZ); National Institutes of Health (NIH) K01DK119375 (KAZ), R01DK115752 (BKY), R01DK129255-01A1 (KAZ), 1R21DK140693- 01 (KAZ), a seed grant from the Presbyterian Health Foundation (PHF; to KAZ), a Wellcome Trust Accelerator Award (314710/Z/24/Z) (DJJ) and a Wellcome Trust Investigator Award (220895/Z/20/Z) (DAL). DJJ is also supported by the Specialised Foundation Programme at the East of England NHS Deanery and DAL’s laboratory is further supported by a NIHR Biomedical Research Centre at Great Ormond Street Hospital for Children NHS Foundation Trust and University College London. The following NIH-funded cores provided services for this project: UAB-UCSD O’Brien Center for Acute Kidney Injury Research P30-DK079337, the UAB Comprehensive Flow Cytometry Core P30-AR048311 and P30-AI27667, and a grant to the Oklahoma Medical Research Foundation (OMRF) core facility (1S10OD028479-01) for purchase of the Aurora Cytek.

---

**Supplemental figure 1. Heatmaps, grouped by metadata variables, showing genes that were significantly different (adjusted p value < 0.05) in the lower limb LOH from control and PKD samples.** Heatmap of DEGs in lower limb LOH between control and PKD samples based on **(A)** sex, **(B)** PKD model, **(C)** day harvested, **(D)** rate of PKD progression, and **(E)** genetic mutation.

**Supplemental figure 2. Heatmaps, grouped by metadata variables, showing genes that were significantly different (adjusted p value < 0.05) in the thin descending limb LOH from control and PKD samples.** Heatmap of DEGs in thin descending limb LOH between control and PKD samples based on **(A)** sex, **(B)** PKD model, **(C)** day harvested, **(D)** rate of PKD progression, and **(E)** genetic mutation.

**Supplemental figure 3. Heatmaps, grouped by metadata variables, showing genes that were significantly different (adjusted p value < 0.05) in principle cells from control and PKD samples.** Heatmap of DEGs in principle cells between control and PKD samples based on **(A)** sex, **(B)** PKD model, **(C)** day harvested, **(D)** rate of PKD progression, and **(E)** genetic mutation.

**Supplemental figure 4. Technology-specific effects on cluster abundance in the proximal tubule. (A)** UMAP showing tubular epithelial cell clustering based on the technology (single cell vs single nucleus). **(B)** Quantification of proximal tubule cluster abundance in each model by technology.

**Supplemental figure 5. Genes increased or decreased in the lower limb LOH based on the type of genetic mutation. (A,B)** Heatmap showing genes that were significantly **(A)** increased or **(B)** decreased in the lower limb LOH of orthologous or non-orthologous mice in relation to non-cystic controls. **(C,D)** Violin plot showing genes that were **(C)** increased or **(D)** decreased in lower limb LOH of both models in relation to non-cystic controls. **(E,F)** Violin plot showing genes that were **(E)** increased or **(F)** decreased in lower limb LOH of both models in relation to non-cystic controls at the sample level.

**Supplemental figure 6. Genes increased or decreased in the lower limb LOH based on the rate of PKD progression. (A,B)** Heatmap showing genes that were significantly **(A)** increased or **(B)** decreased in the lower limb LOH (in relation to non-cystic controls) based on the rate of PKD progression. **(C,D)** Violin plot showing genes that were **(C)** increased or **(D)** decreased in lower limb LOH of both models in relation to non-cystic controls. **(E,F)** Violin plot showing genes that were **(E)** increased or **(F)** decreased in lower limb LOH of both models in relation to non-cystic controls at the sample level.

**Supplemental figure 7. Examples of model specific changes in gene expression at the single cell level. (A,B)** *Igfbp7* expression at the **(A)** group or **(B)** individual sample level. **(C,D)** *Tmem27* expression at the **(C)** group or **(D)** sample level. **(E)** *Lcn2* expression in individual models.

**Supplemental figure 8. Upset plot and Jaccard indices of genes increased or decreased in mixed S1/S2, S2 proximal tubule, or S1 proximal tubule in relation to non-cystic controls from each model. (A-C)** Upset plot and Jaccard indices of genes increased or decreased in relation to non-cystic controls in the **(A)** Mixed S1/S2 **(B)** S2 proximal tubule, or **(C)** S1 proximal tubule. Genes highlighted in red are increased in all slow PKD models; genes highlighted in blue are increased in all rapid PKD models; genes highlighted in green are enriched in all non-orthologous models; genes highlighted in black were identified in pseudobulk analysis.

**Supplemental figure 9. Upset plot and Jaccard indices of genes increased or decreased in the thick ascending limb LOH, S3 proximal tubule, and distal convoluted tubule in relation to non-cystic controls from each model. (A-C)** Upset plot and Jaccard indices of genes increased or decreased in relation to non-cystic controls in the **(A)** thick ascending limb LOH **(B)** S3 proximal tubule, or **(C)** distal convoluted tubule. Genes highlighted in red are increased in all slow PKD models; genes highlighted in blue are increased in all rapid PKD models; genes highlighted in green are enriched in all non-orthologous models; genes highlighted in black were identified in pseudobulk analysis.

**Supplemental figure 10. Upset plot and Jaccard indices of genes increased or decreased in the nephron connecting tubule, intercalated cells, and nephron connecting tubule 2 in relation to non-cystic controls from each model. (A-C)** Upset plot and Jaccard indices of genes increased or decreased in relation to non-cystic controls in the **(A)** nephron connecting tubule **(B)** intercalated cells, or **(C)** nephron connecting tubule 2. Genes highlighted in red are increased in all slow PKD models; genes highlighted in blue are increased in all rapid PKD models; genes highlighted in green are enriched in all non-orthologous models; genes highlighted in black were identified in pseudobulk analysis.

**Supplemental figure 11. scRNAseq atlas of mouse PKD.** UMAP showing the fully annotated scRNAseq atlas of mouse PKD.

**Supplemental figure 12. Signaling interactions between PKD-enriched cluster and all other cell types at the whole atlas and individual model level. (A-G)** Number and strength of interactions between PKD enriched clusters and all other cell types in the **(A)** combined atlas and **(B-G)** each individual model. Red indicates that the communication between the two cell types is increased in PKD while blue means it is decreased in PKD.

**Supplemental figure 13. Spatial transcriptomics. (A)** H&E stained images of control samples from *Pkd1*^WT/WT^ and *Ift88* mice.

**Supplemental figure 14. TACCO deconvolution of control *Ift88* sample.** Deconvolution of all cell types shown in Supplemental figure 11 in control *Ift88* sample.

**Supplemental figure 15. TACCO deconvolution of control *Ift88* sample.** Deconvolution of all cell types shown in Supplemental figure 11 in control *Ift88* sample.

**Supplemental figure 16. TACCO deconvolution of *Ift88* sample.** Deconvolution of all cell types shown in Supplemental figure 11 in *Ift88* sample.

**Supplemental figure 17. TACCO deconvolution of *Ift88* sample.** Deconvolution of all cell types shown in Supplemental figure 11 in *Ift88* sample.

**Supplemental figure 18. TACCO deconvolution of control *Pkd1*^WT/WT^ sample.**

Deconvolution of all cell types shown in Supplemental figure 11 in control *Pkd1*^WT/WT^ sample.

**Supplemental figure 19. TACCO deconvolution of control *Pkd1*^WT/WT^ sample.**

Deconvolution of all cell types shown in Supplemental figure 11 in control *Pkd1*^WT/WT^ sample.

**Supplemental figure 20. TACCO deconvolution of *Pkd1*^RC/RC^ sample.** Deconvolution of all cell types shown in Supplemental figure 11 in *Pkd1*^RC/RC^ sample.

**Supplemental figure 21. TACCO deconvolution of *Pkd1*^RC/RC^ sample.** Deconvolution of all cell types shown in Supplemental figure 11 in *Pkd1*^RC/RC^ sample.

**Supplemental figure 22. Violin plot showing *Spp1* expression in all spots of control or *Ift88*mice.**

**Supplemental figure 23. Ligand receptor interactions in cystic niches of *Ift88* and *Pkd1*^RC/RC^ mice. (A)** Outlined cystic and non-cystic niches used for ligand-receptor analysis. **(B, C)** Top 50 ligand-receptor interactions between PKD enriched clusters and all other cell types in the kidney in **(B)** *Ift88* and **(C)** *Pkd1*^RC/RC^ mice. Arrows indicate ligand-receptor pairs that were also present in scRNAseq data from each respective model.

**Supplemental figure 24. Figure 6. Cyst size does not impact gene expression in PKD. (A)** Outline of cystic regions binned based on cyst size (large, medium, small). **(B)** Principle component analysis (PCA) of top 100 highly variable genes (HVGs) in each sized cyst from *Pkd1*^RC/RC^ or *Ift88* mice. **(C)** DEGs in large vs small cysts of *Pkd1*^RC/RC^ or *Ift88* mice. **(D)** DEGs in large vs medium cysts of *Pkd1*^RC/RC^ or *Ift88* mice. **(E)** DEGs in medium vs small cysts of *Pkd1*^RC/RC^ or *Ift88* mice.

**Supplemental figure 25. DEGs in different sized cysts in relation to one another and non- cystic spots. (A)** DEGs in large cysts vs normal tissue in *Pkd1*^RC/RC^ or *Ift88* mice. **(B)** DEGs in medium cysts vs normal tissue in *Pkd1*^RC/RC^ or *Ift88* mice. **(C)** DEGs in small cysts vs normal tissue in *Pkd1*^RC/RC^ or *Ift88* mice.

**Supplemental figure 26. Analysis of individual cysts and spots in *Pkd1*^RC/RC^ and *Ift88* mice. (A)** Individual cysts outlined in *Pkd1*^RC/RC^ and *Ift88* mice. **(B)** Top 100 highly variable genes across individual cysts in *Pkd1*^RC/RC^ and *Ift88* mice. **(C)** Top 100 highly variable genes across all spots found in cystic regions in *Pkd1*^RC/RC^ and *Ift88* mice.

**Supplemental figure 27. Spots with injury, inflammation, and fibrosis signature mapped back to tissue sections in *Pkd1*^RC/RC^ and *Ift88* mice.**

**Supplemental figure 28. *Spp1* expression in cystic and non-cystic regions of *Pkd1*^RC/RC^ mice.**

**Supplemental figure 29. Dotplot showing ligand receptor pairs between lower limb PKD cluster 2, thin limb PKD cluster 1, and other cell types in the kidney in scRNAseq data.**

**Supplemental figure 30. Top 50 ligand receptor pairs between lower limb PKD cluster 2, thin limb PKD cluster 1, and other cell types in the kidney from spatial transcriptomics data.**

**Supplemental figure 31. Immune cell numbers in *Pkd1*^RC/RC^ mice with and without *Spp1.*** Flow cytometry quantification of immune cell numbers in control, *Pkd1*^RC/RC^ *Spp1*^WT^, *Pkd1*^RC/RC^ *Spp1*-/- mice harvested at one year of age.

**Supplemental figure 32. Visual step-by-step guide to navigate the single cell website.** A. Home page display includes links to individual datasets, single-cell ATLAS visualization, dataset downloads, as well as help and feedback. The home page also displays dataset details split by species and sex. B. The ATLAS web page features a large UMAP display of selected ATLAS (human or mouse), which can then be filtered by cell type, sex, model (mouse ATLAS only), or disease status. Gene expression can be visualized for selected and filtered ATLAS, and is displayed both overlaid onto a UMAP as well as across cell clusters via violin plot at the bottom of the page.

## Notes

https://bmblx.bmi.osumc.edu/scPKD/

### Competing Interest Statement

The authors have declared no competing interest.

