## Supplemental Figures 1-32 for "A cross model spatial and single-cell atlas reveals the conserved involvement of osteopontin in polycystic kidney disease"

### Supplemental figure 1

#### Lower limb LOH

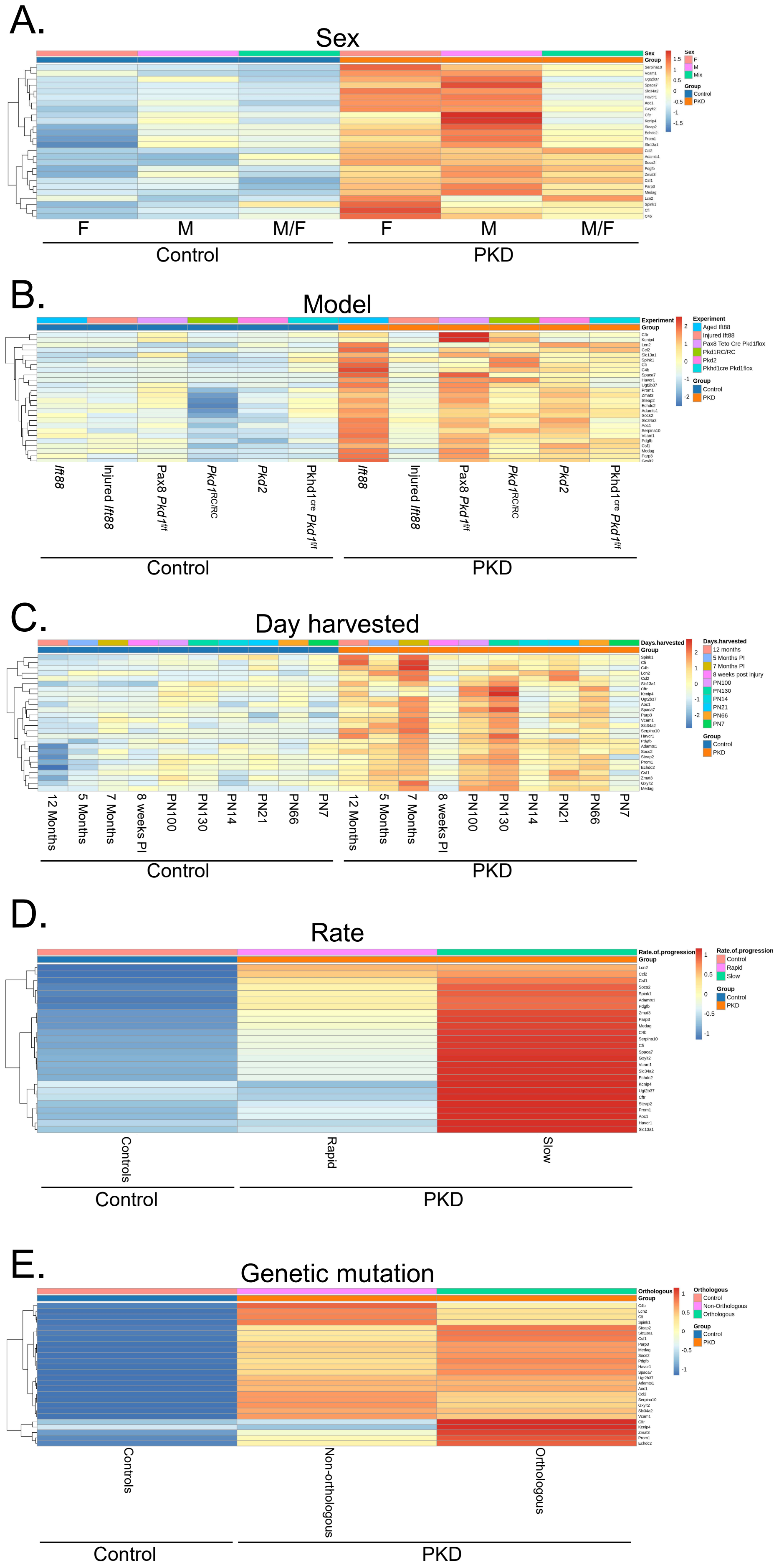

### Supplemental figure 2

#### Thin descending limb LOH

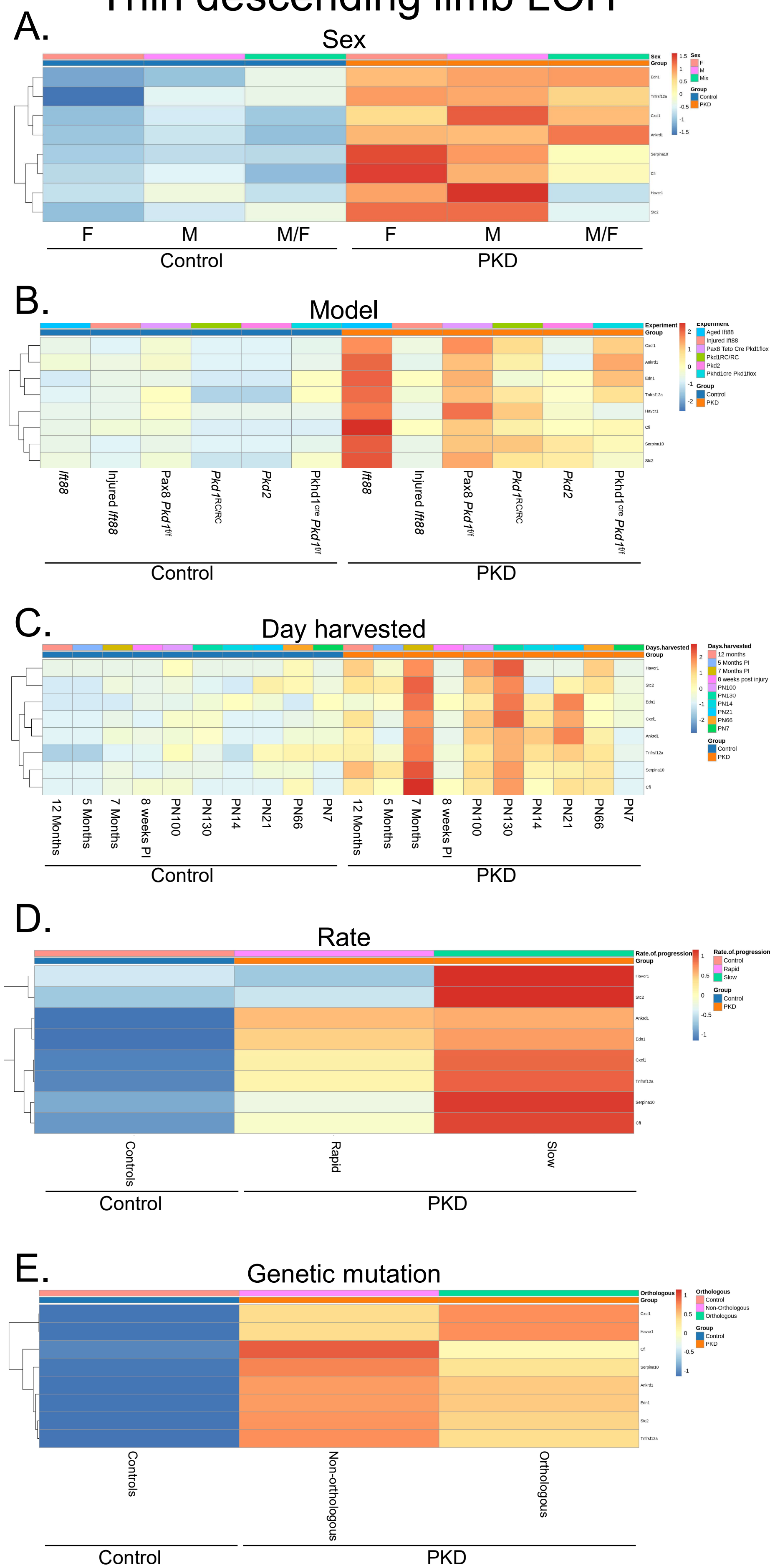

### Supplemental figure 3

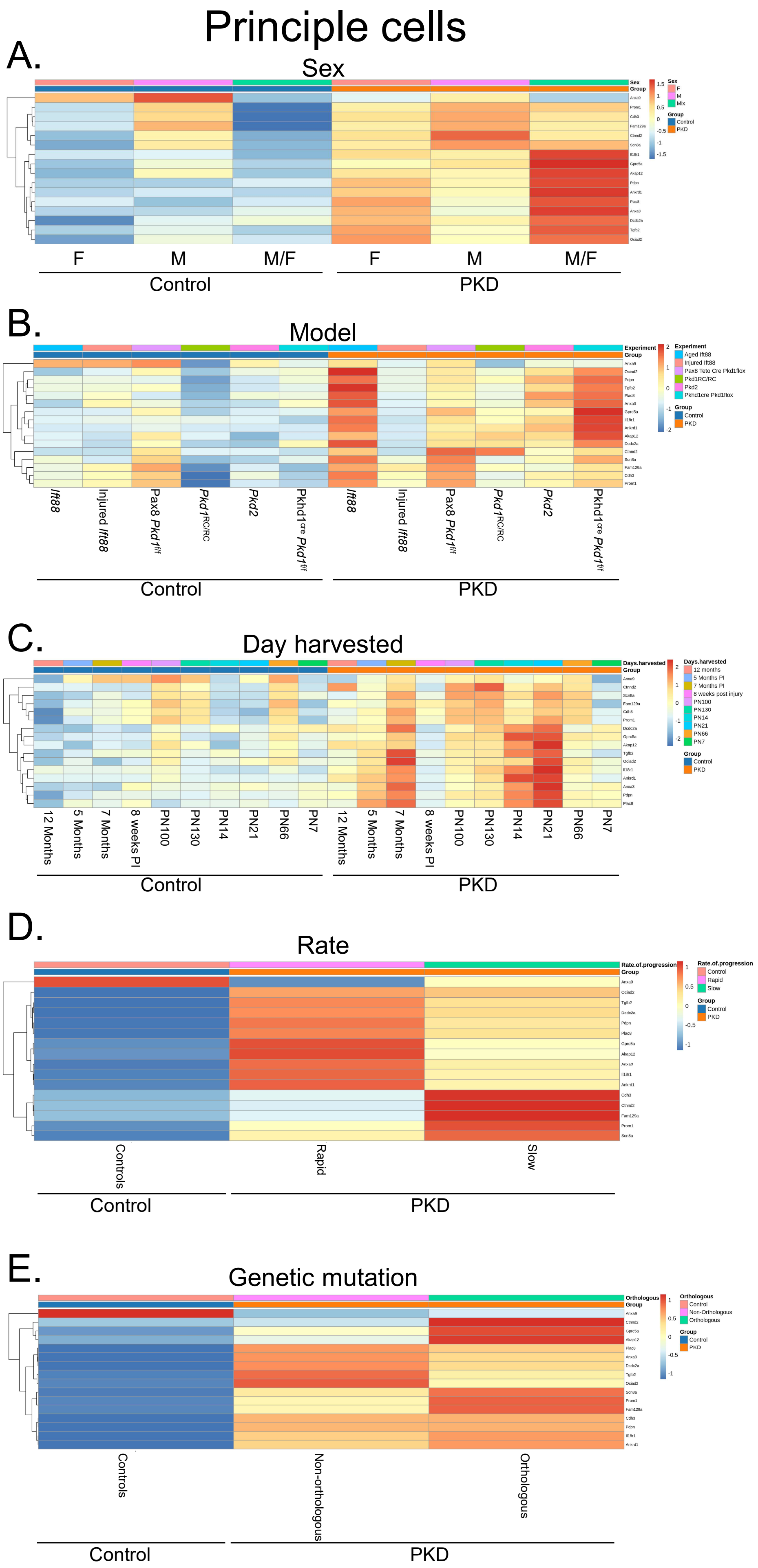

### Supplemental figure 4

A.

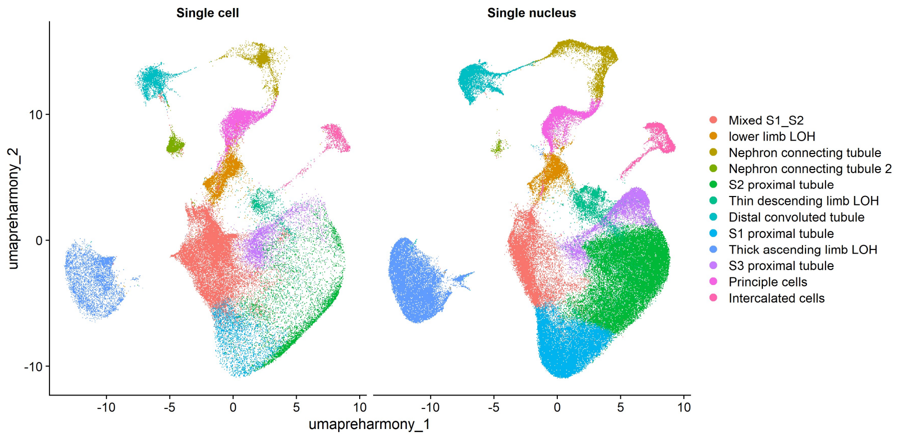


B.

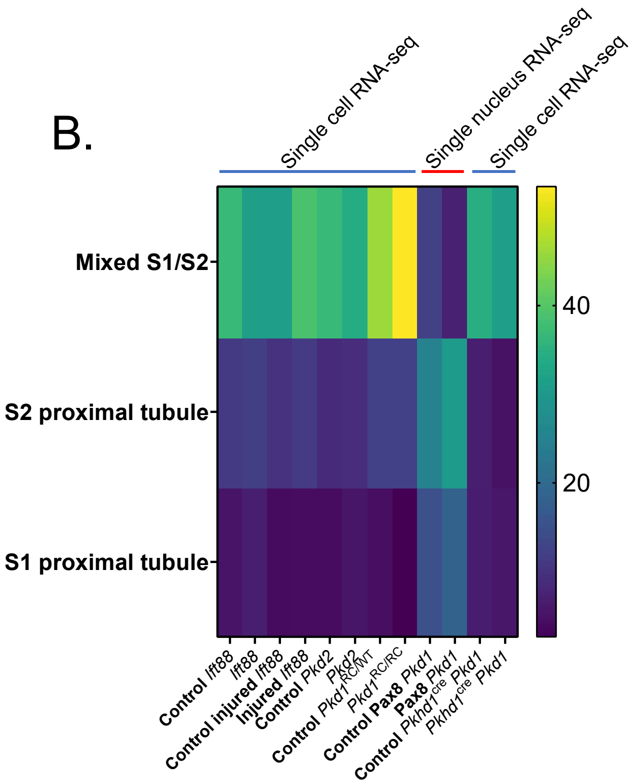

### Supplemental figure 5

#### Lower limb LOH

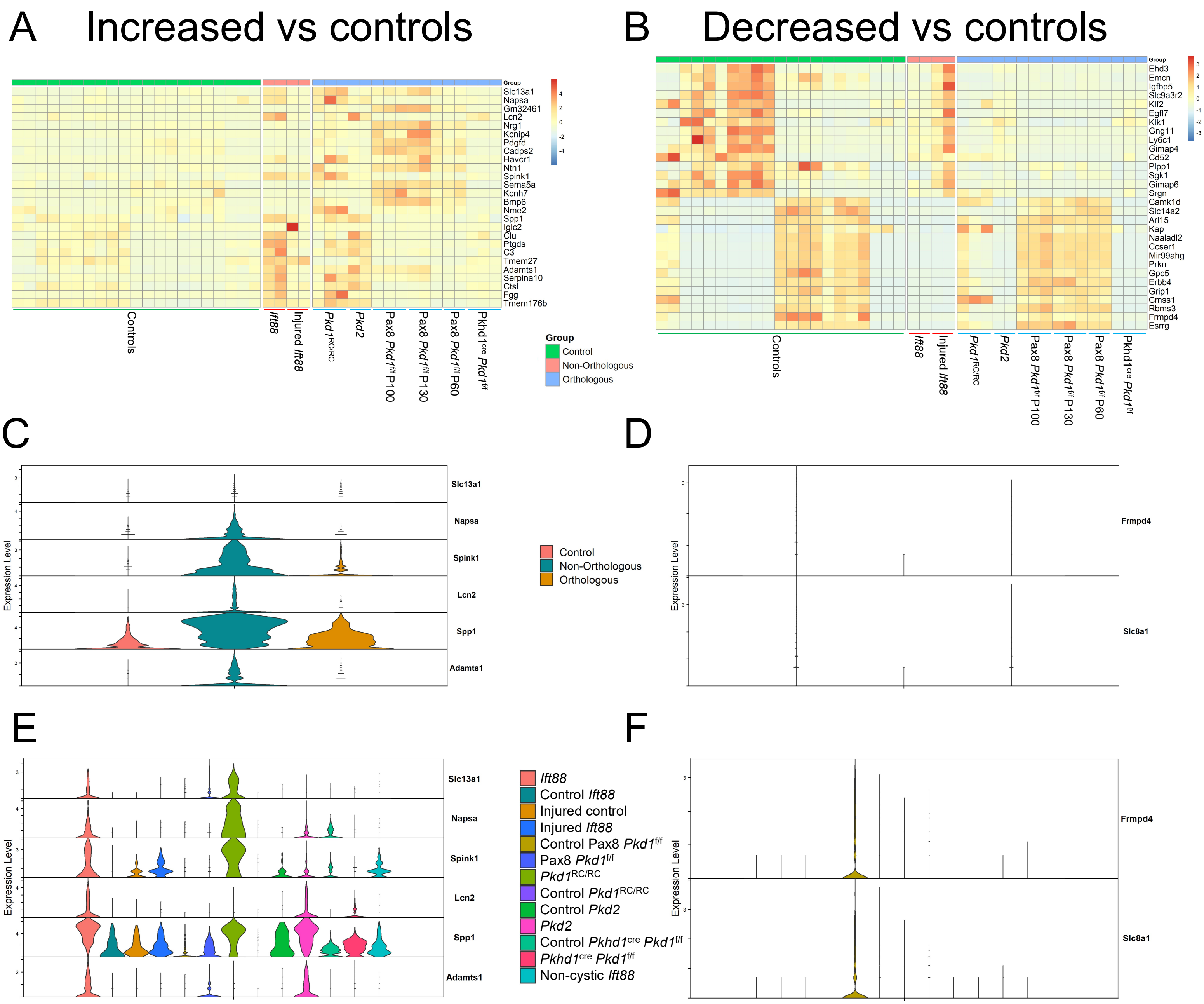

### Supplemental figure 6

#### Lower limb LOH

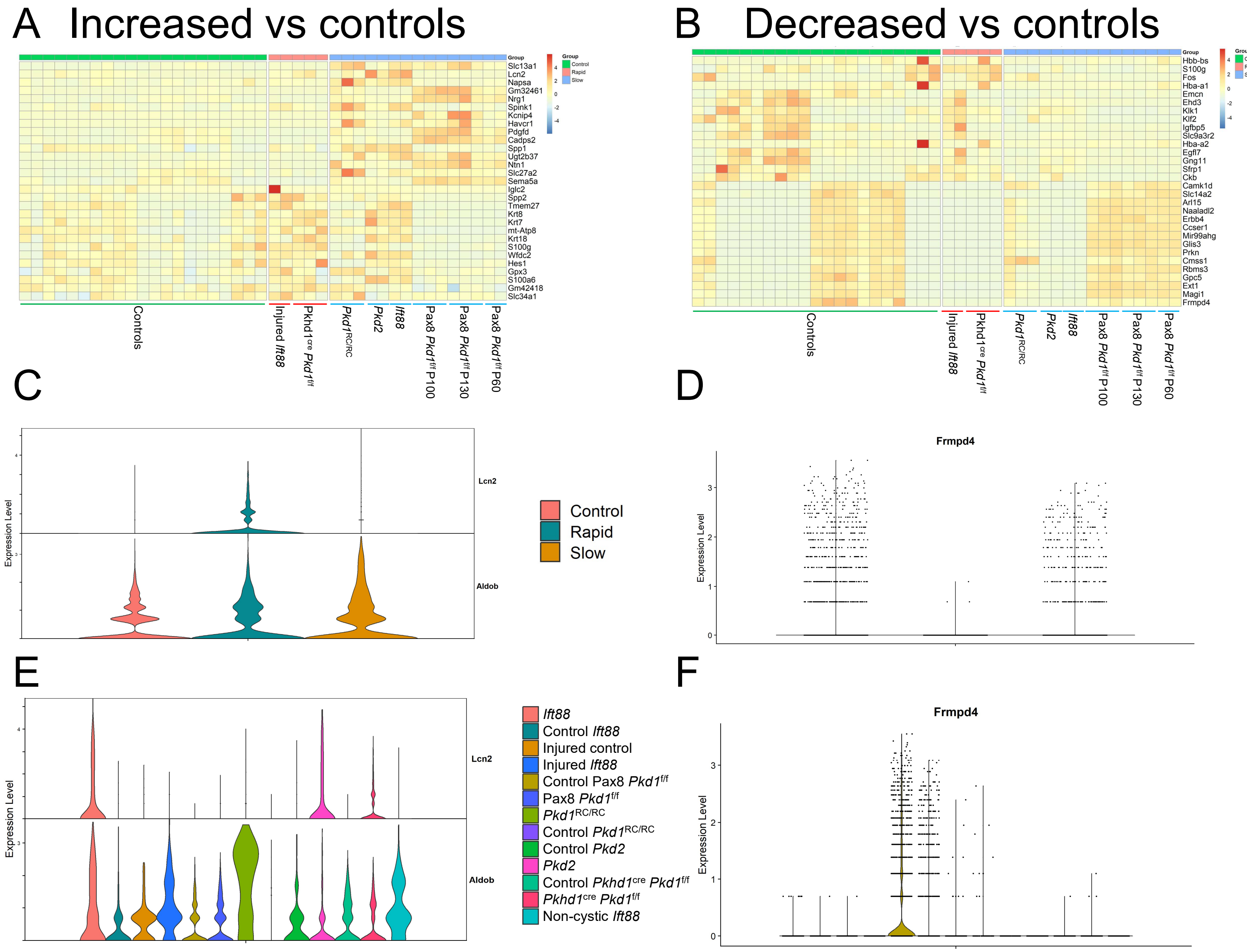

### Supplemental figure 7

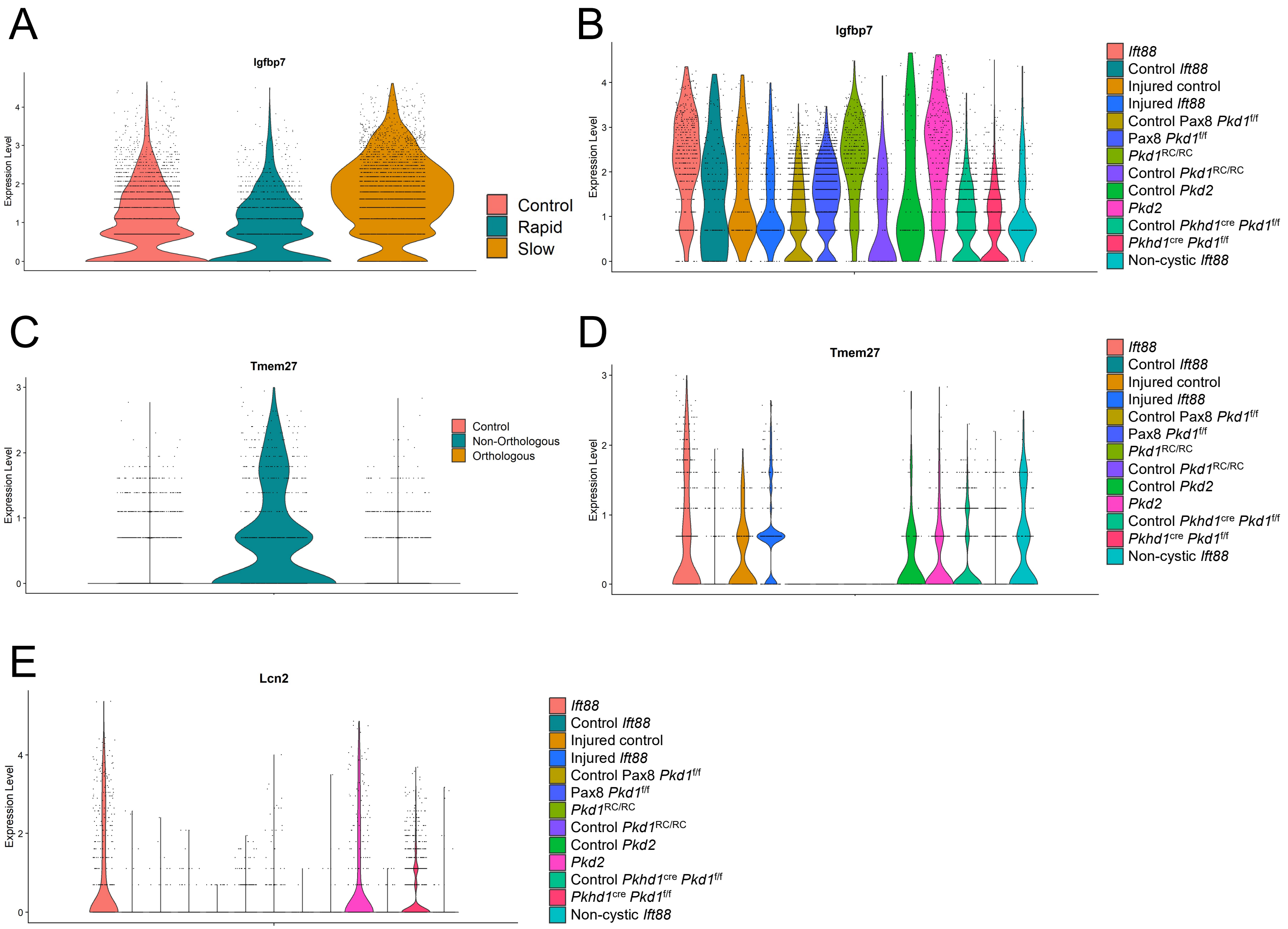

### Supplemental figure 8

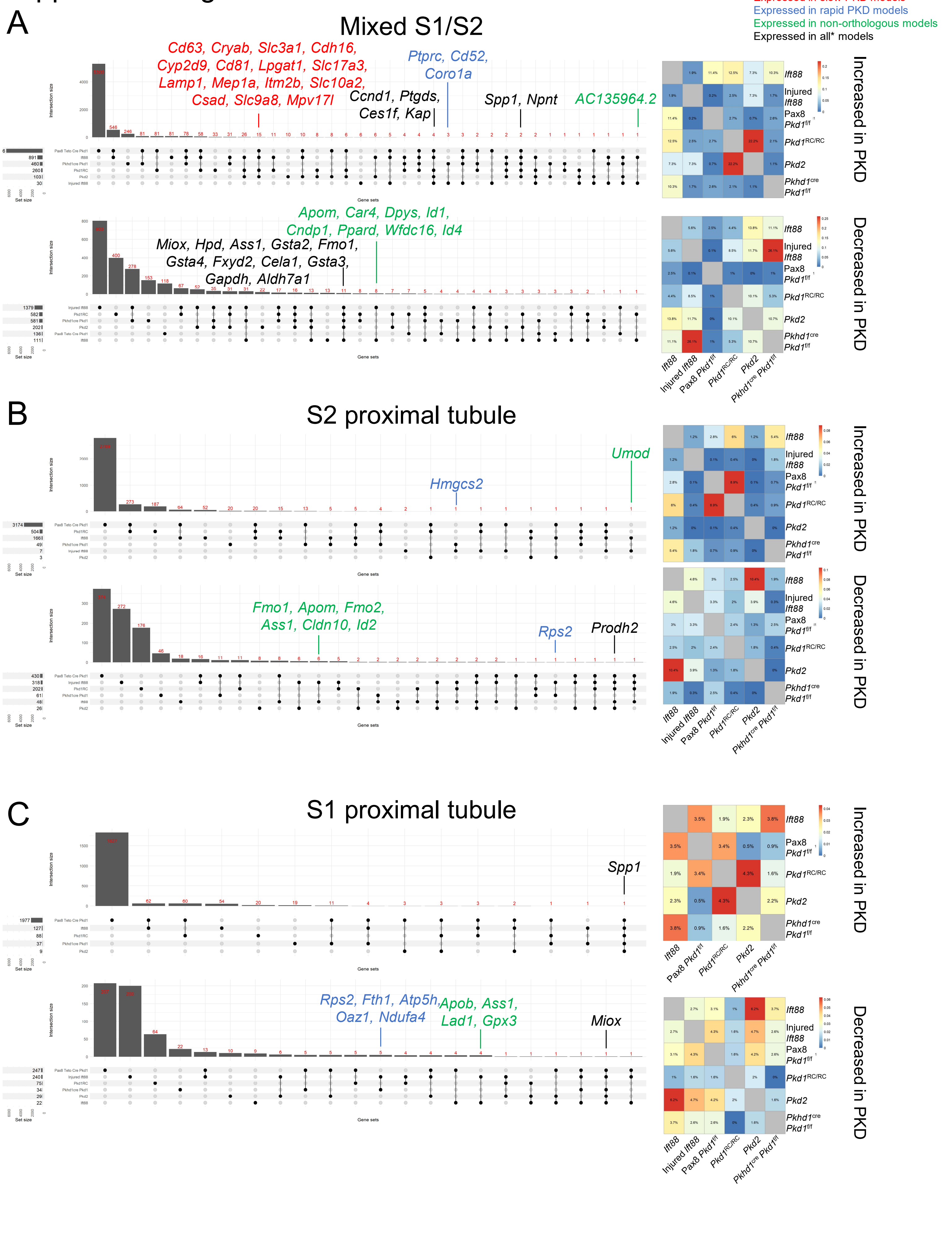

### Supplemental figure 9

Expressed in slow PKD models
Expressed in rapid PKD models
Expressed in orthologous models
Expressed in non-orthologous models
Expressed in all* models

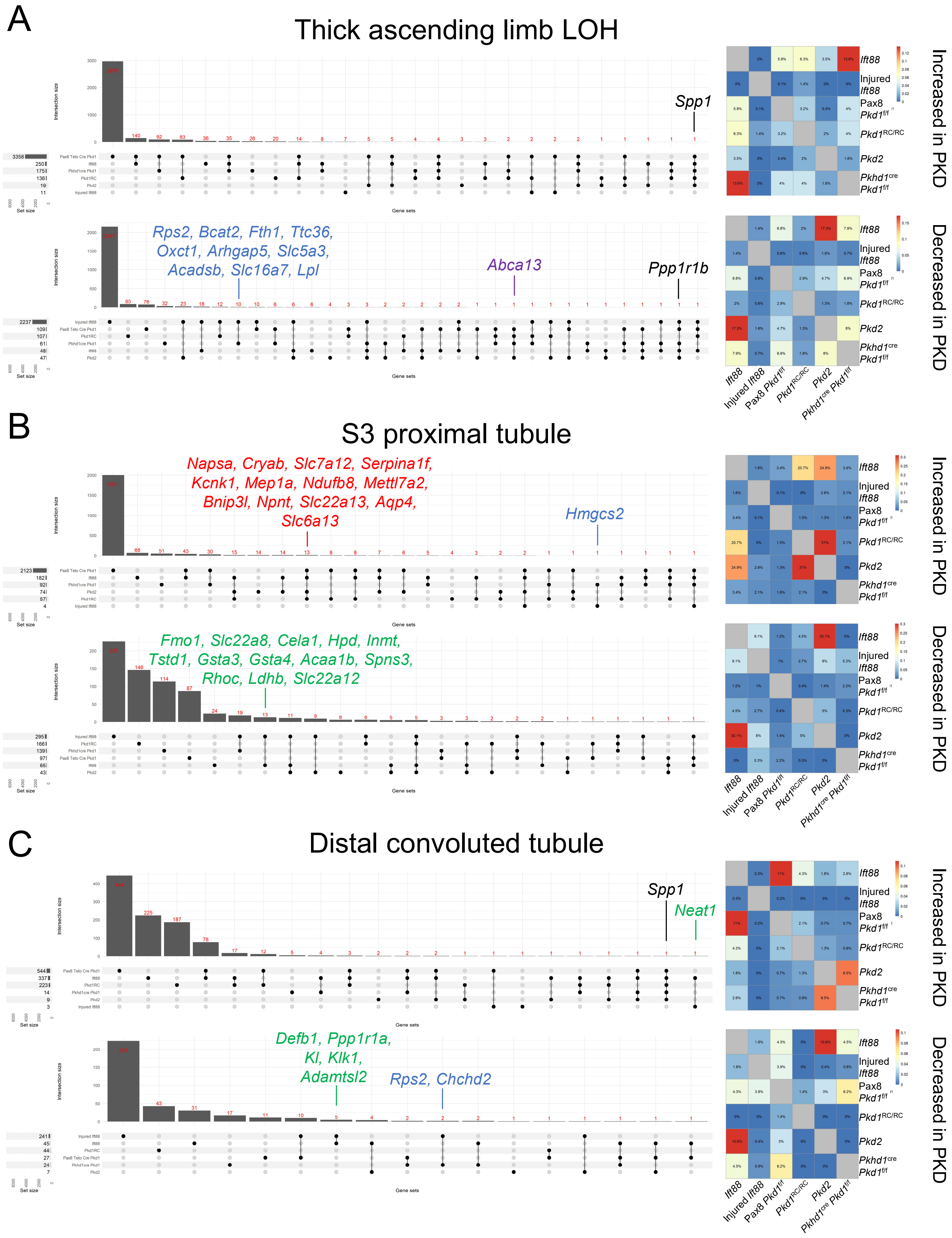

### Supplemental figure 10

Expressed in slow PKD models
Expressed in rapid PKD models
Expressed in non-orthologous models
Expressed in all* models

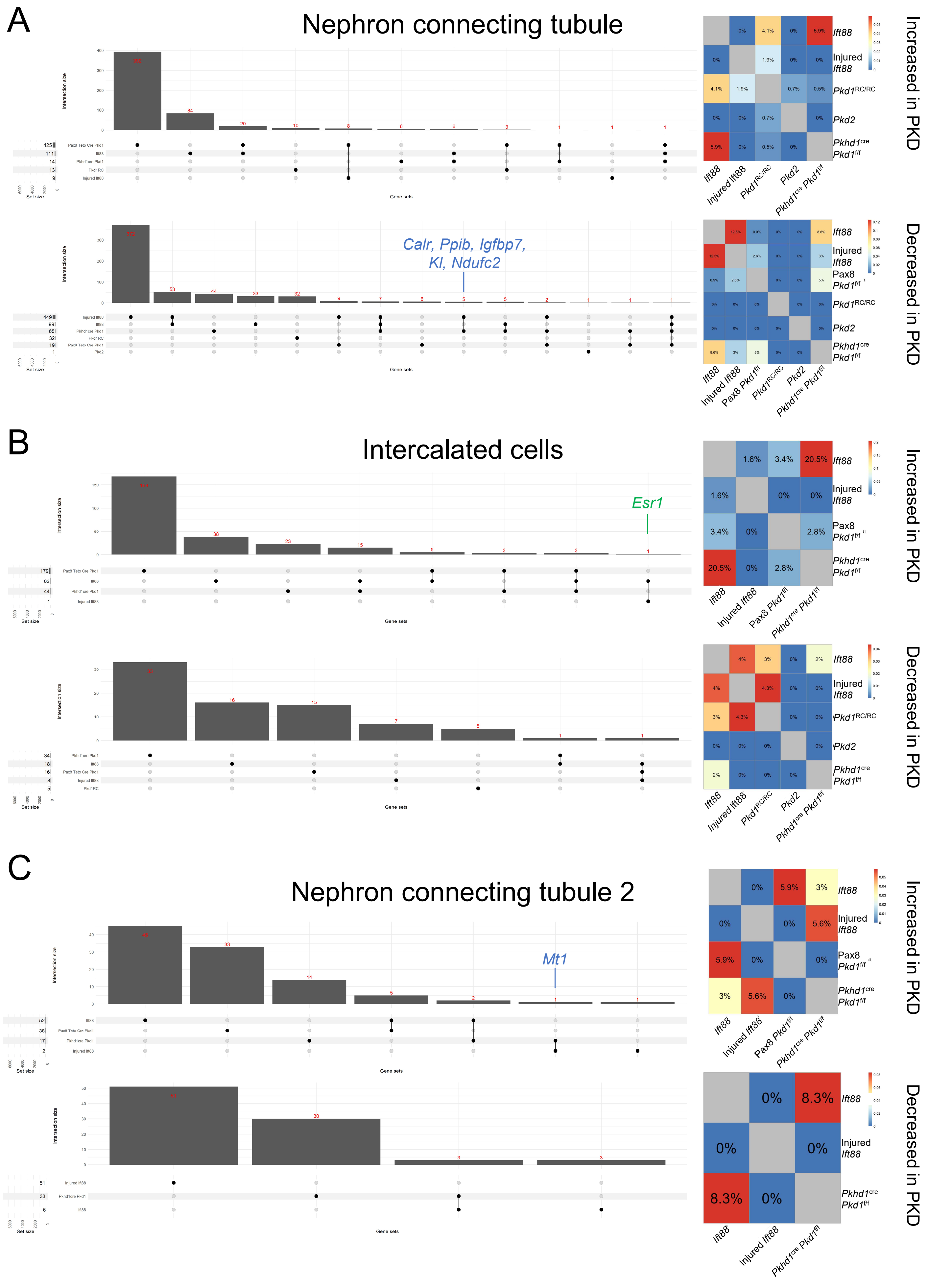

### Supplemental figure 11

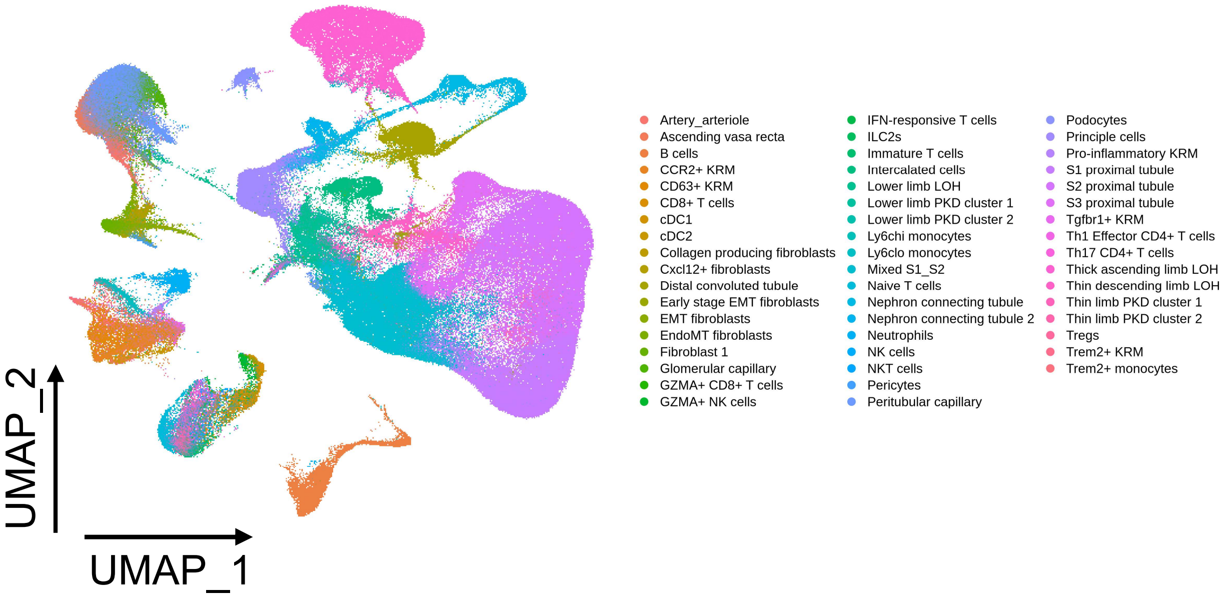

### Supplemental figure 12

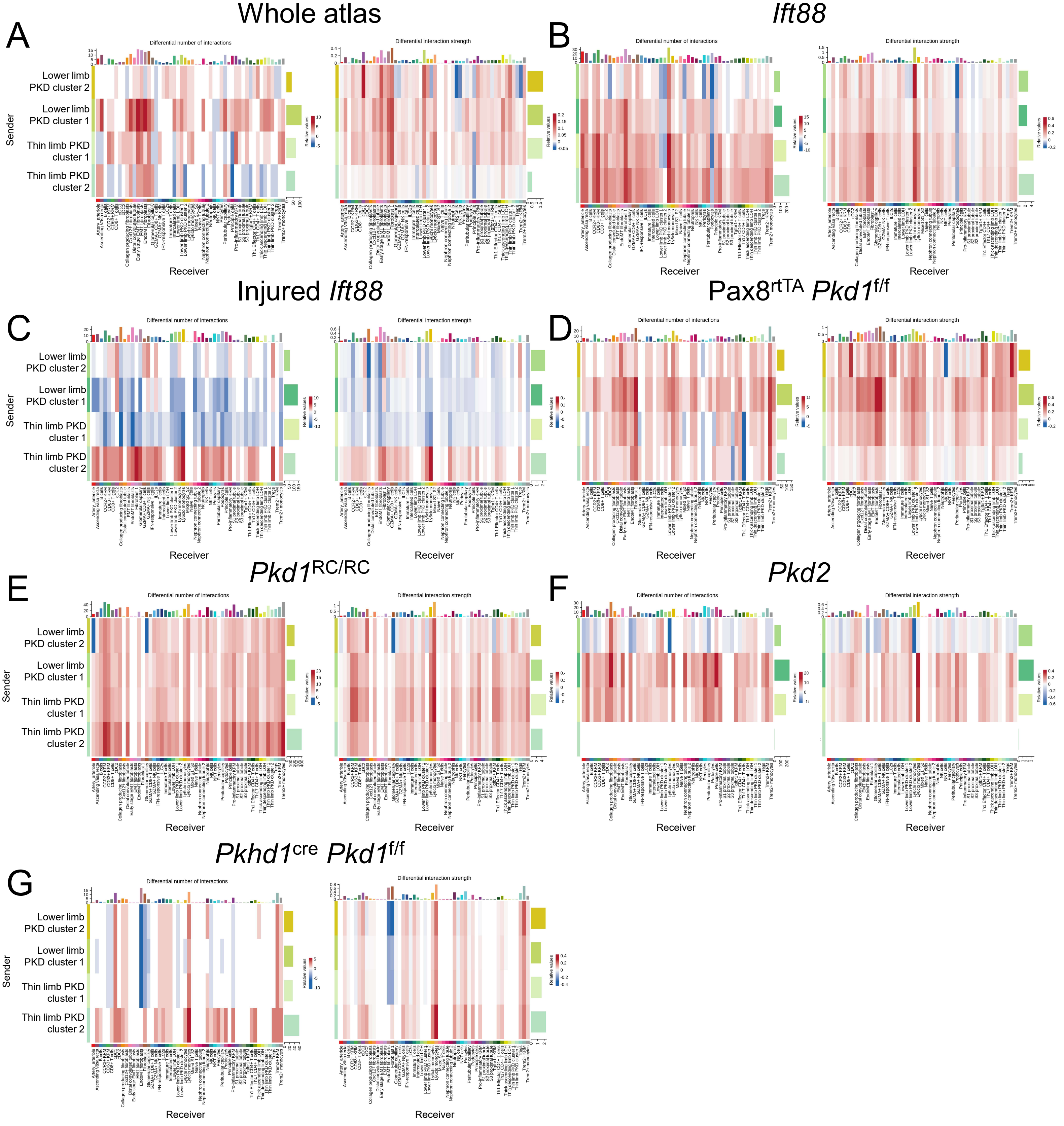

### Supplemental figure 13

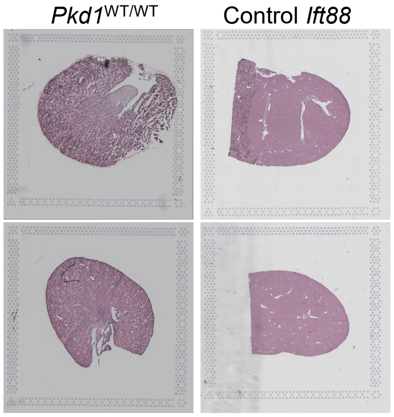

### Supplemental figure 14

Control *Ift88*

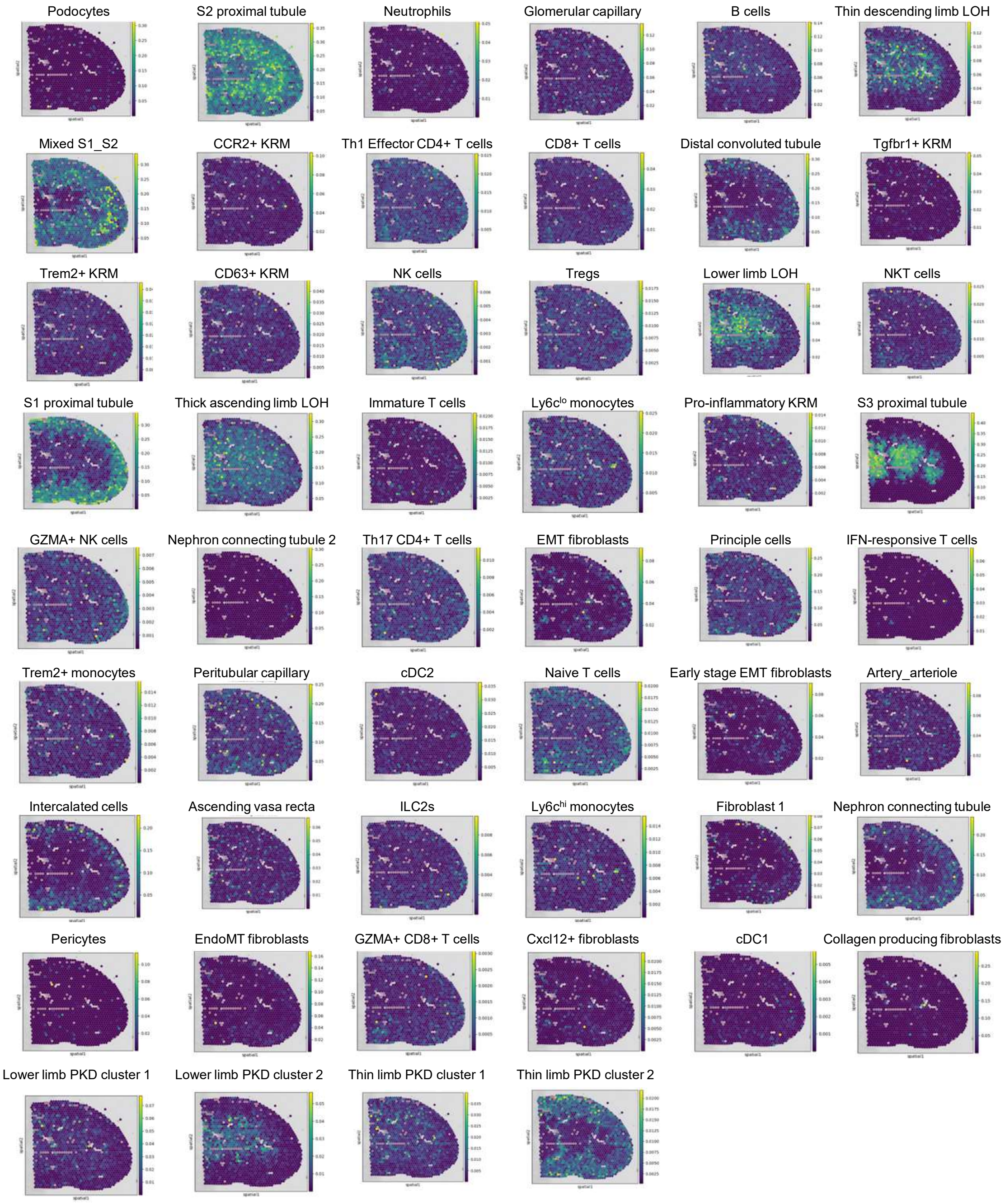

### Supplemental figure 15

#### Control *Ift88*

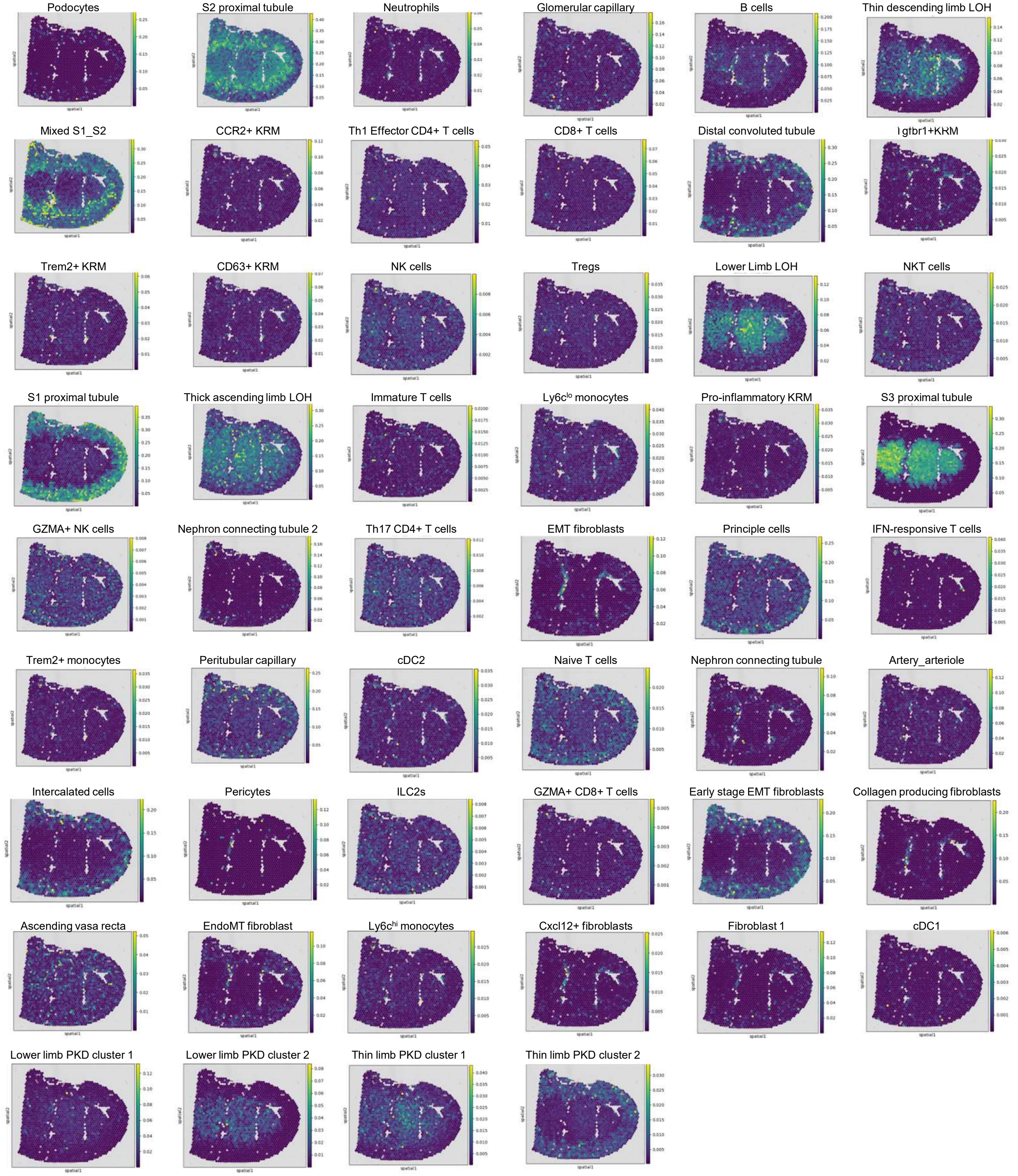

### Supplemental figure 16

*Ift88*

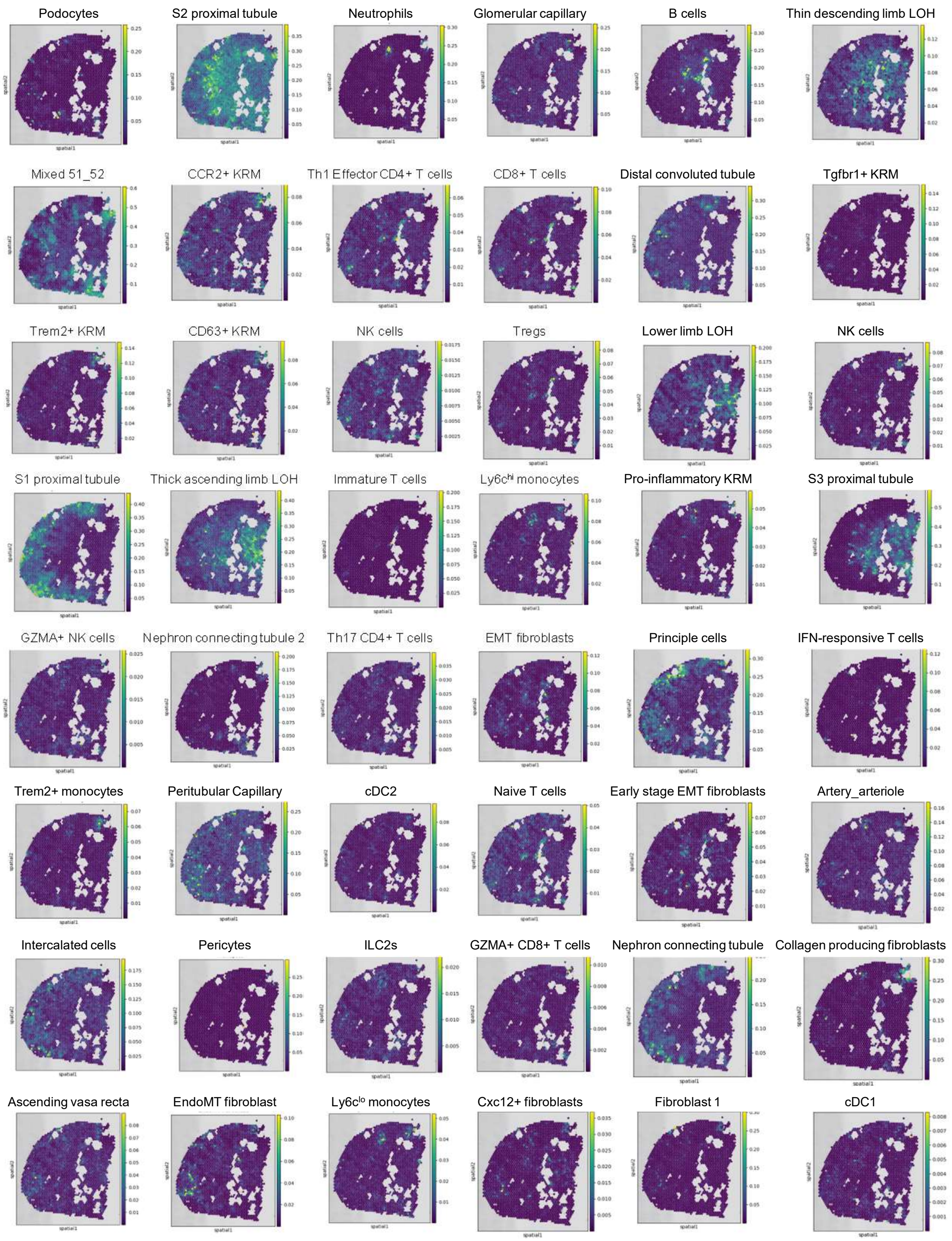

### Supplemental figure 17

*Ift88*

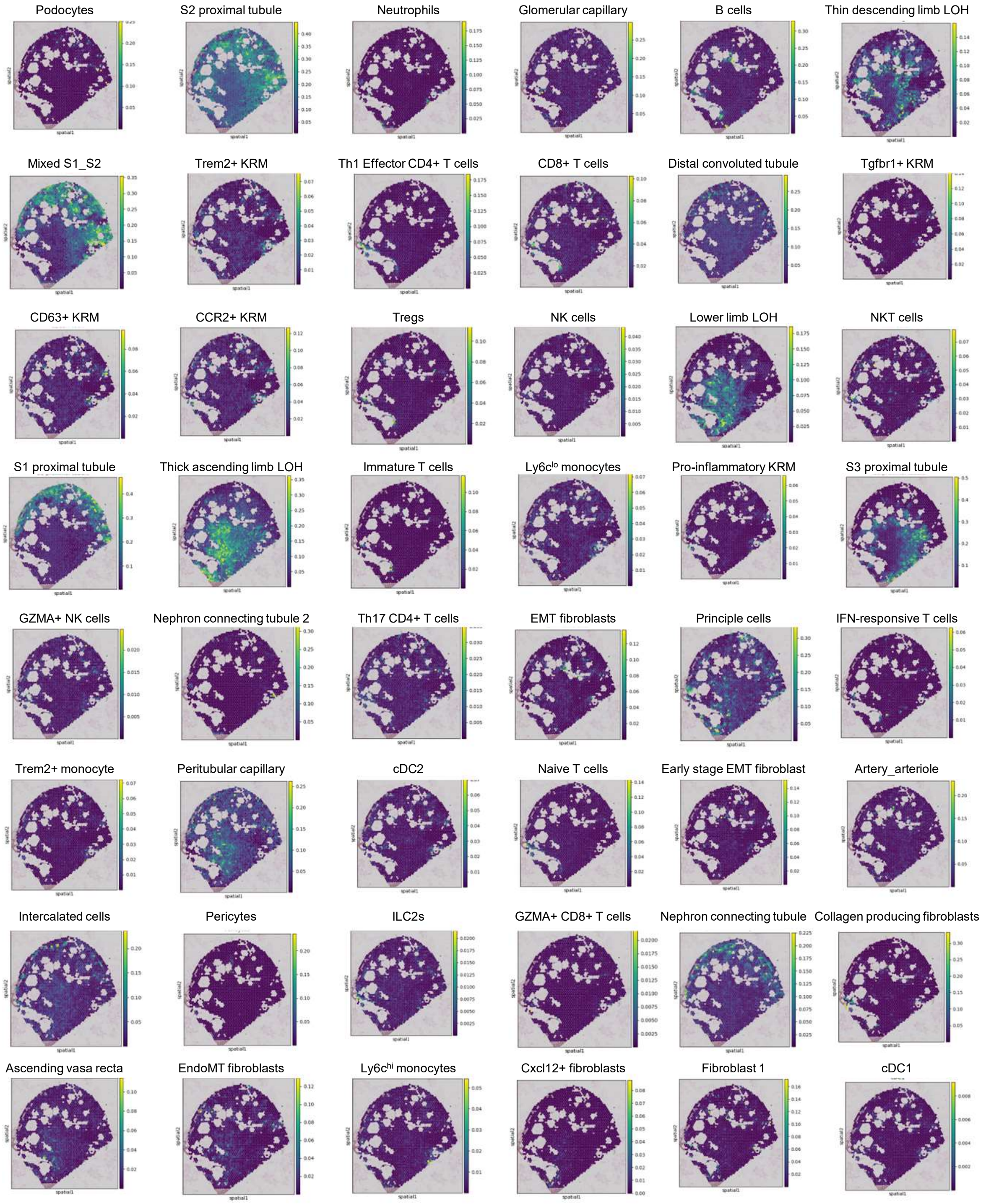

### Supplemental figure 18

#### Control *Pkd1*$^{WT/WT}$

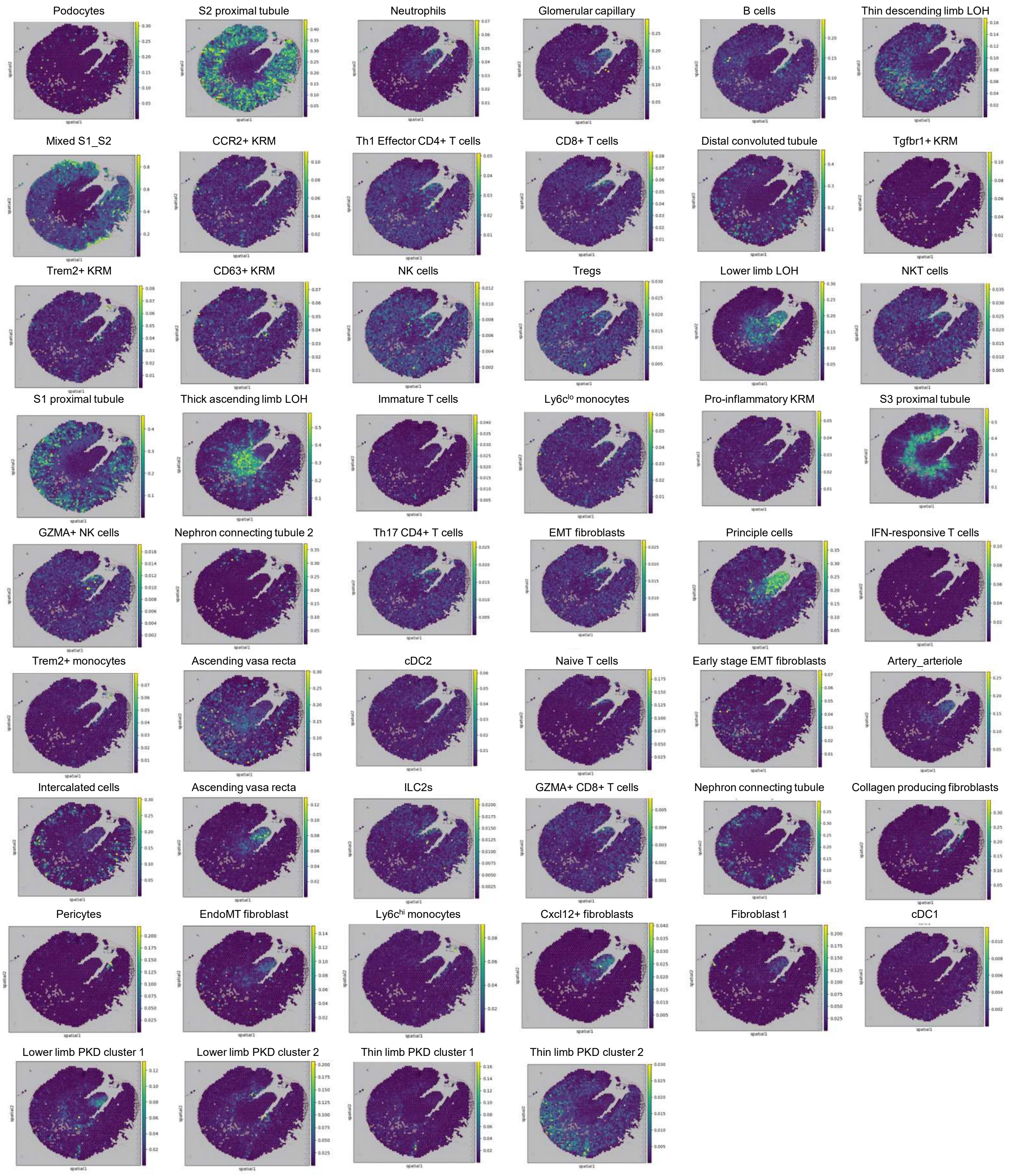

### Supplemental figure 19

#### Control *Pkd1*$^{WT/WT}$

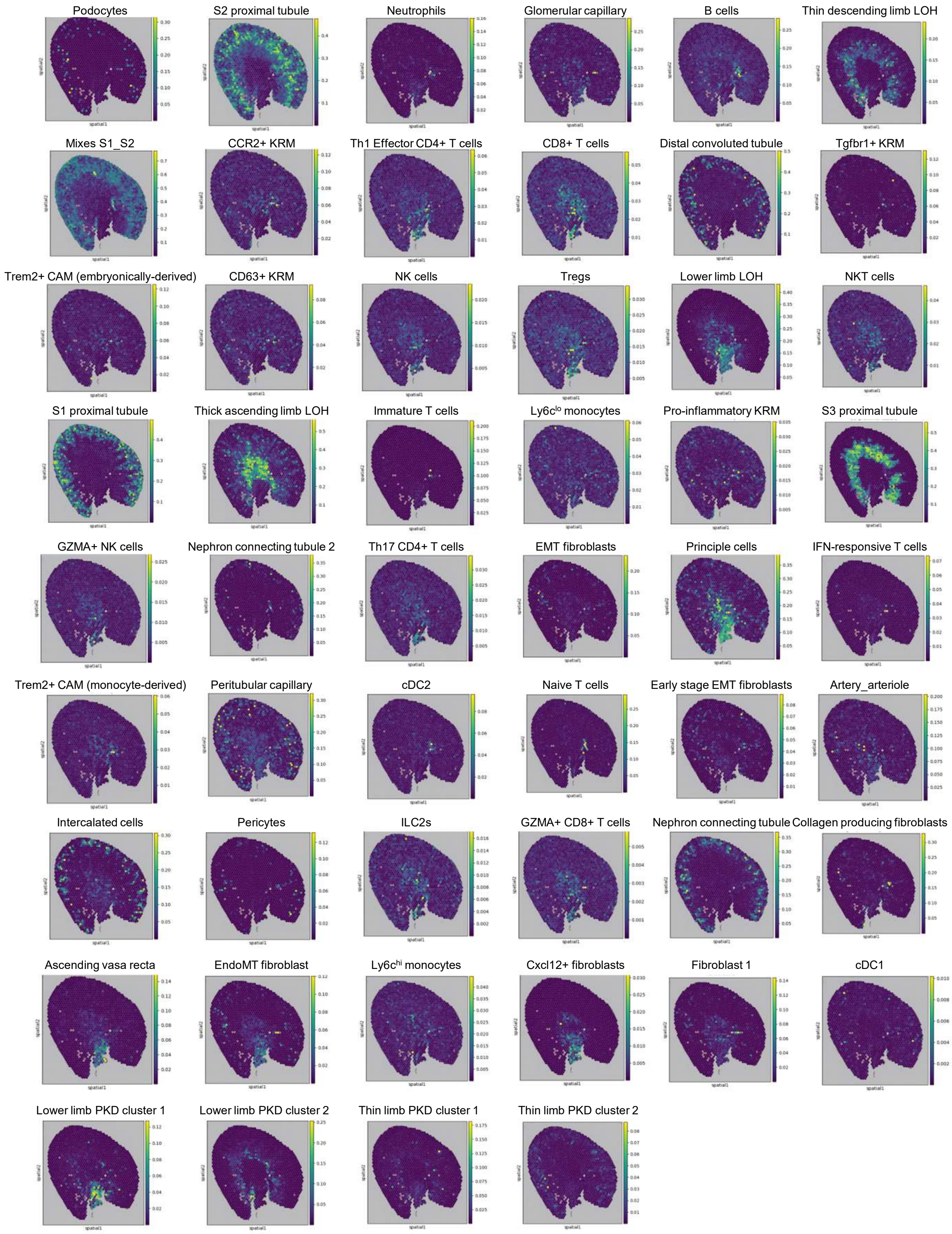

### Supplemental figure 20

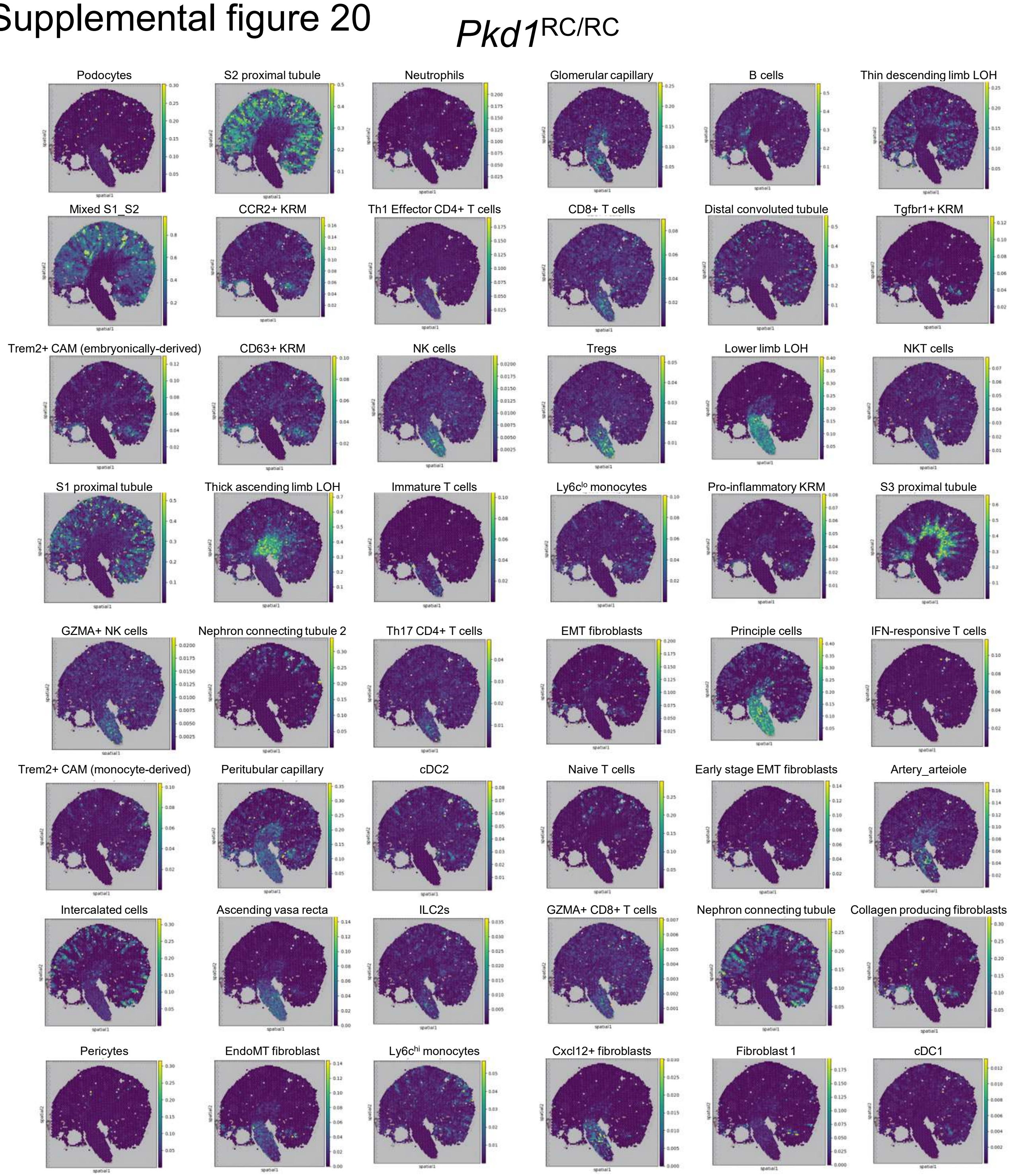

### Supplemental figure 21

*Pkd1*$^{RC/RC}$

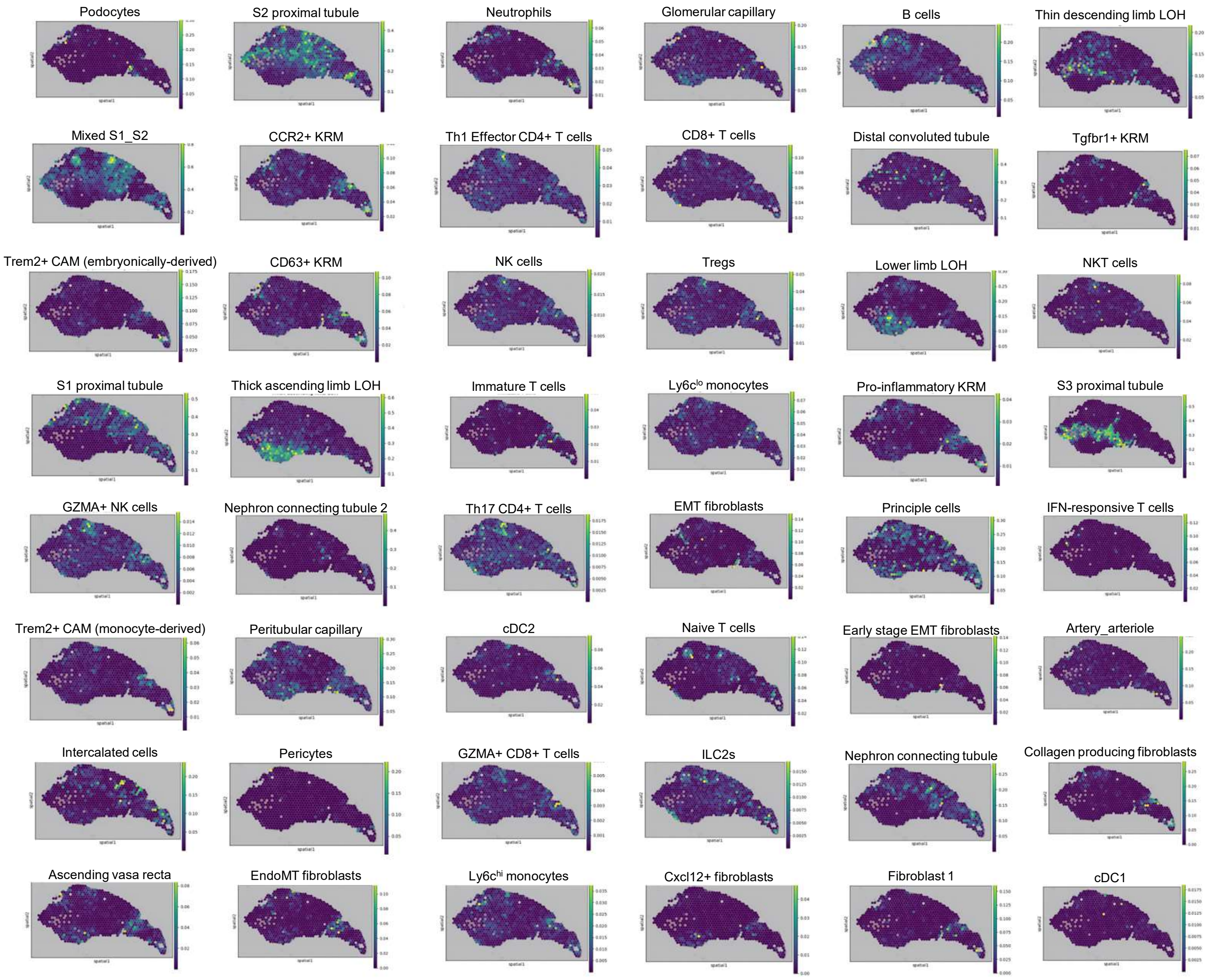

Supplemental figure 22

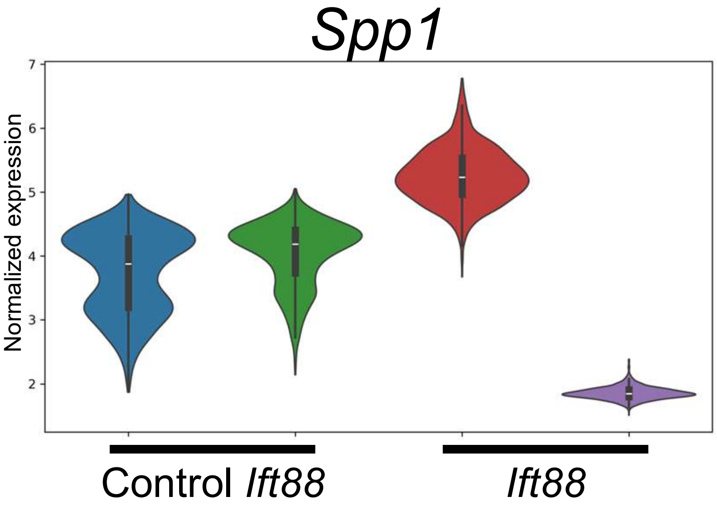

### Supplemental figure 23

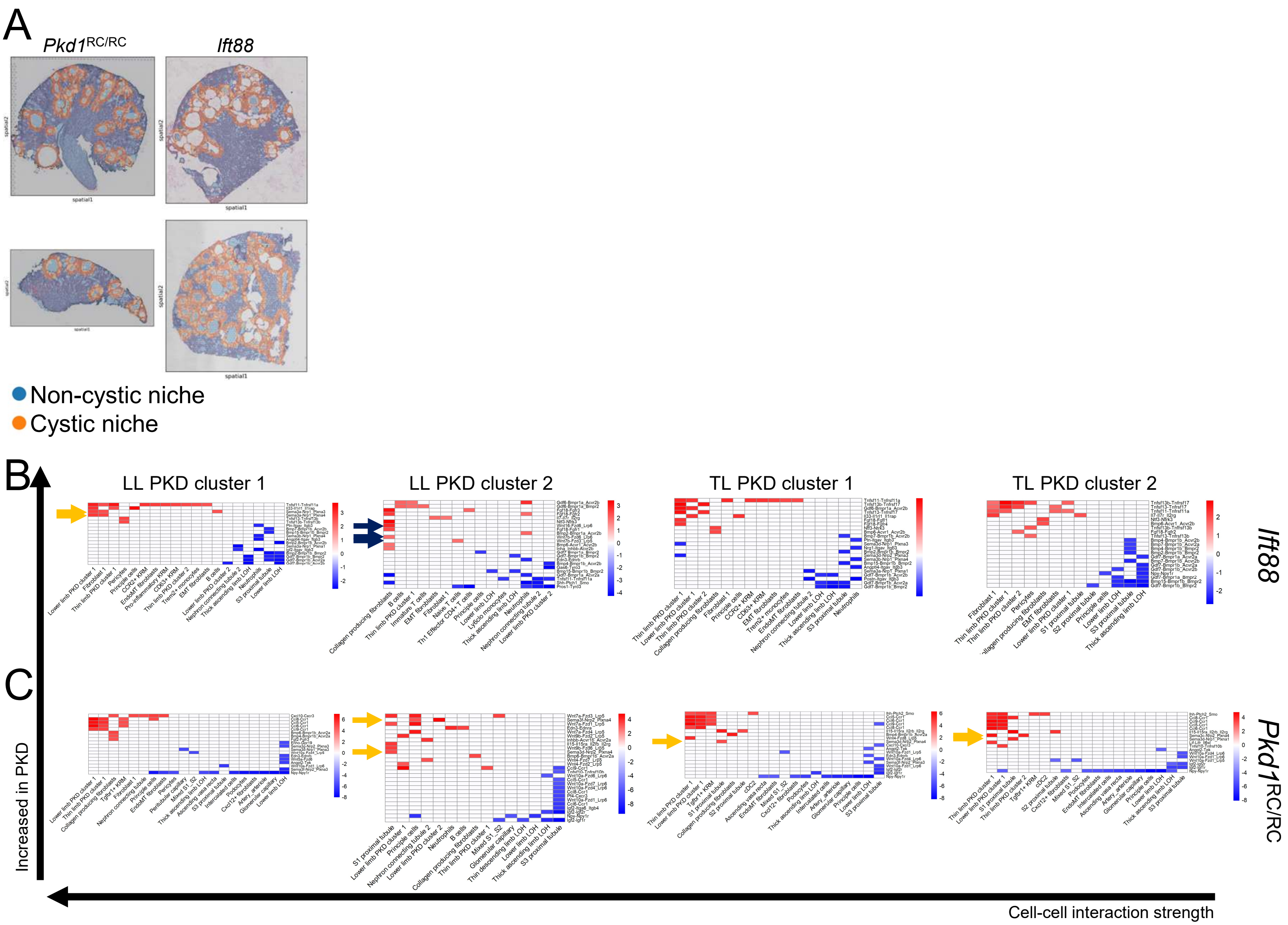

Supplemental figure 24

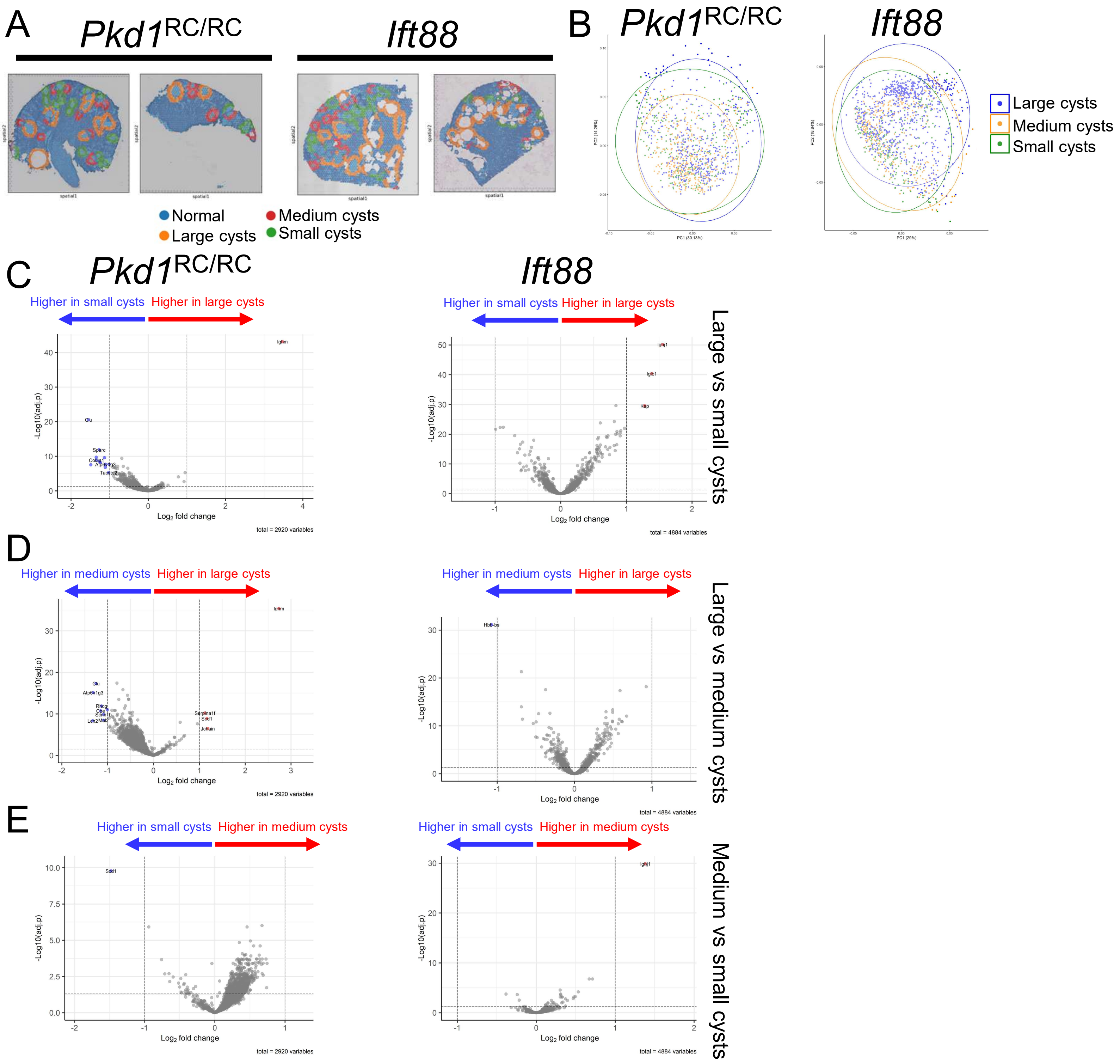

### Supplemental figure 25

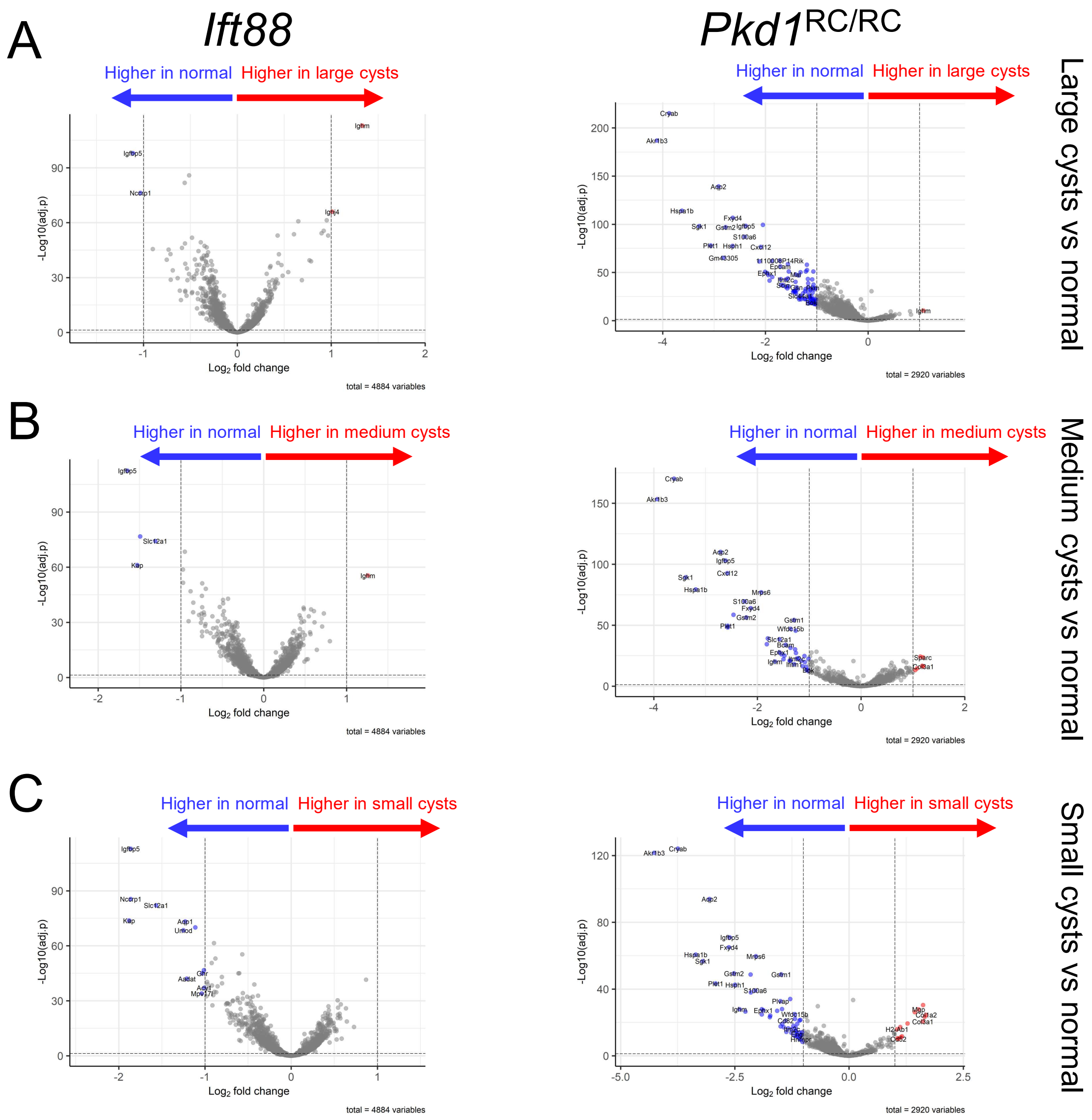

Supplemental figure 26

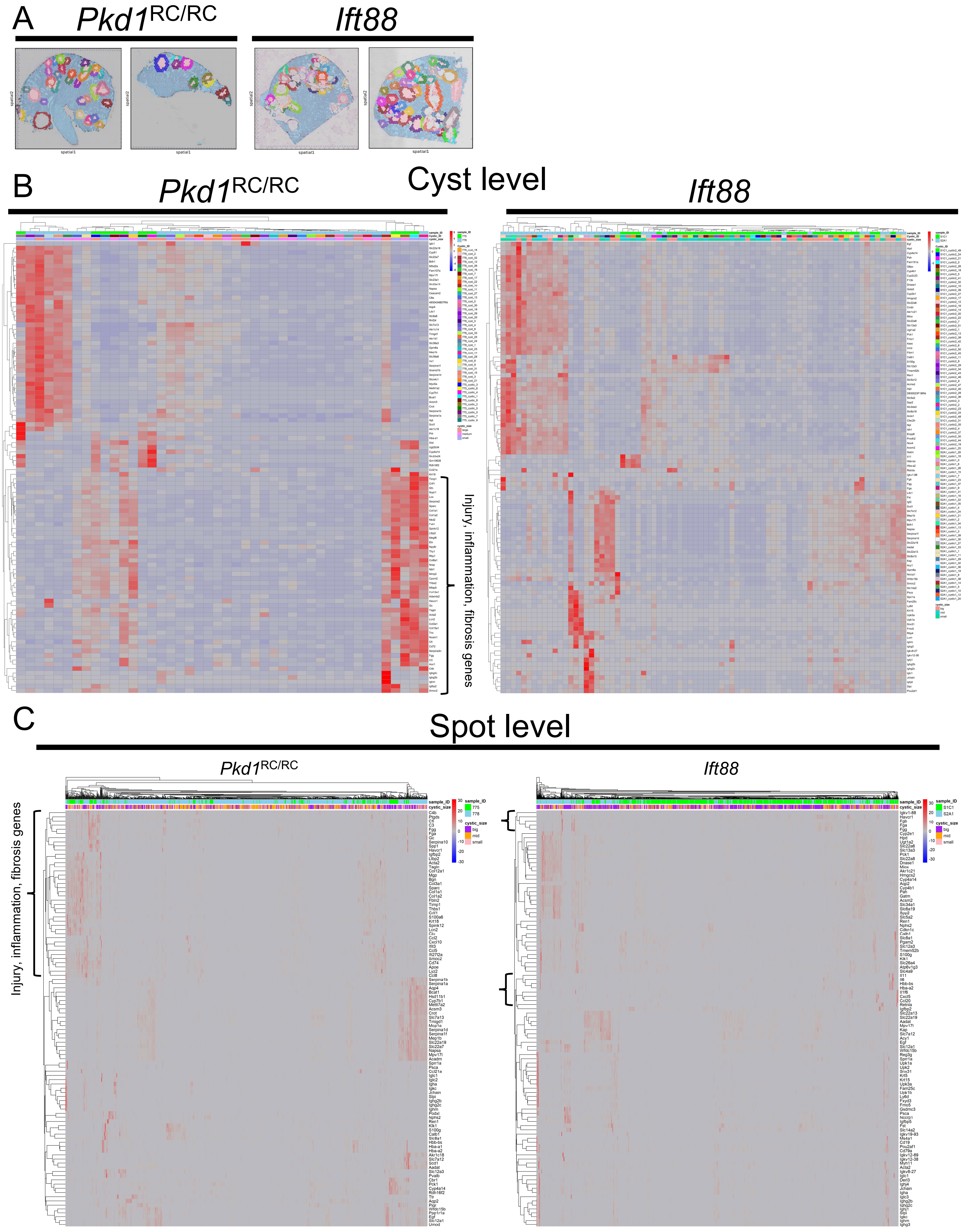

### Supplemental figure 27

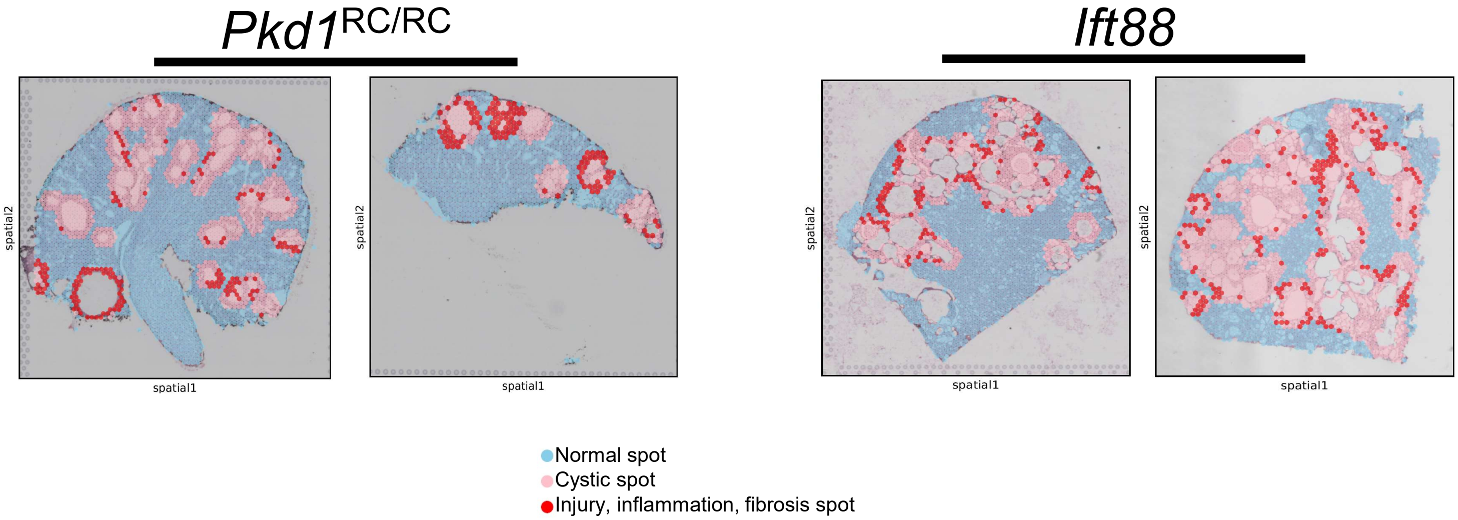

### Supplemental figure 28

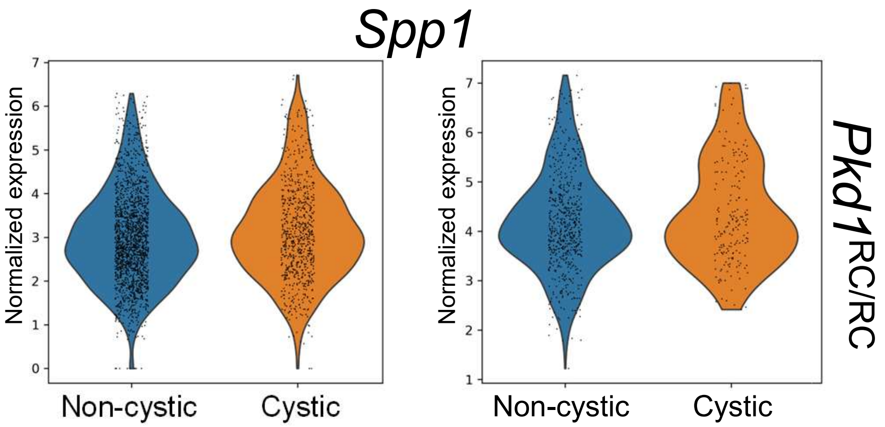

### Supplemental figure 29

#### Lower limb PKD cluster 2

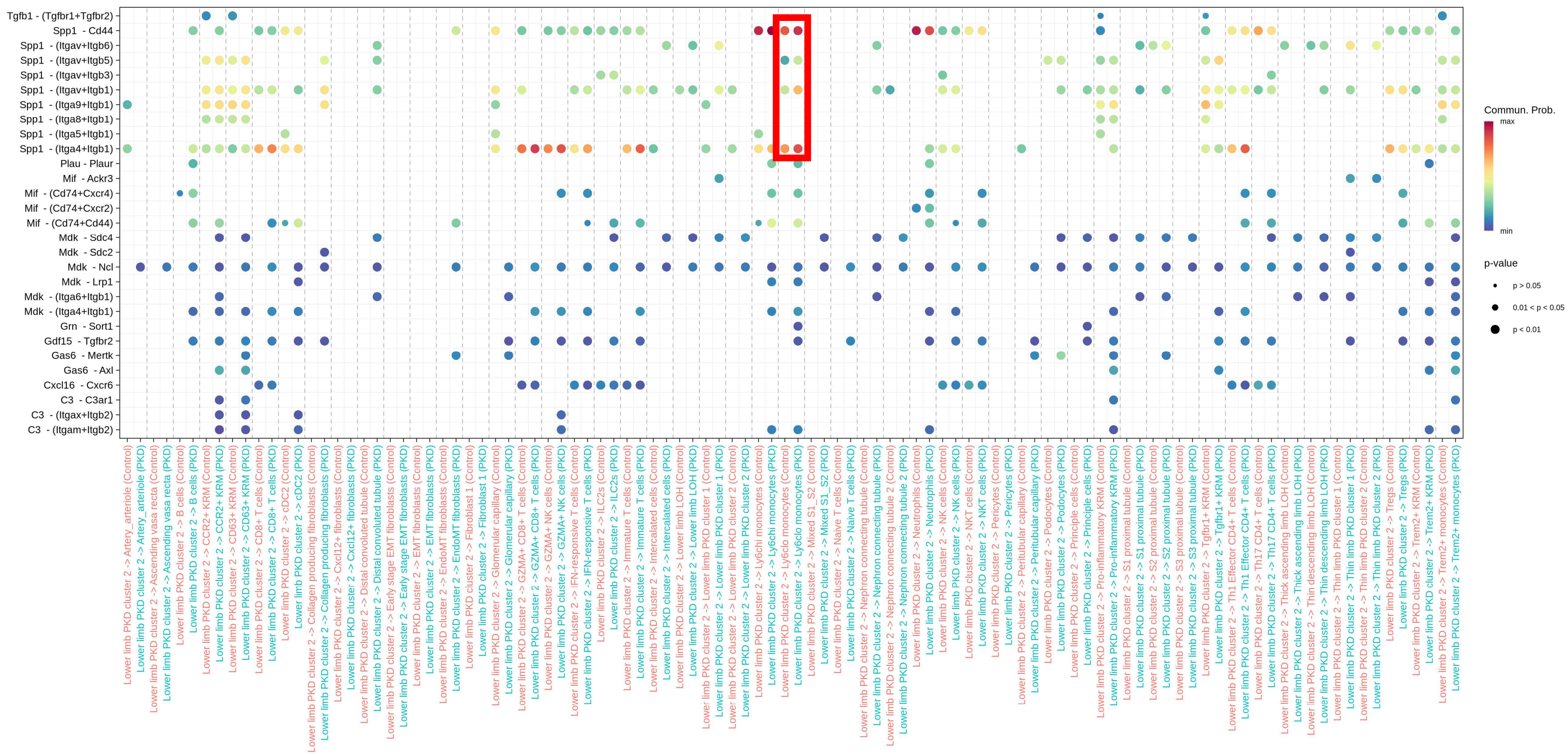


#### Thin limb PKD cluster 1

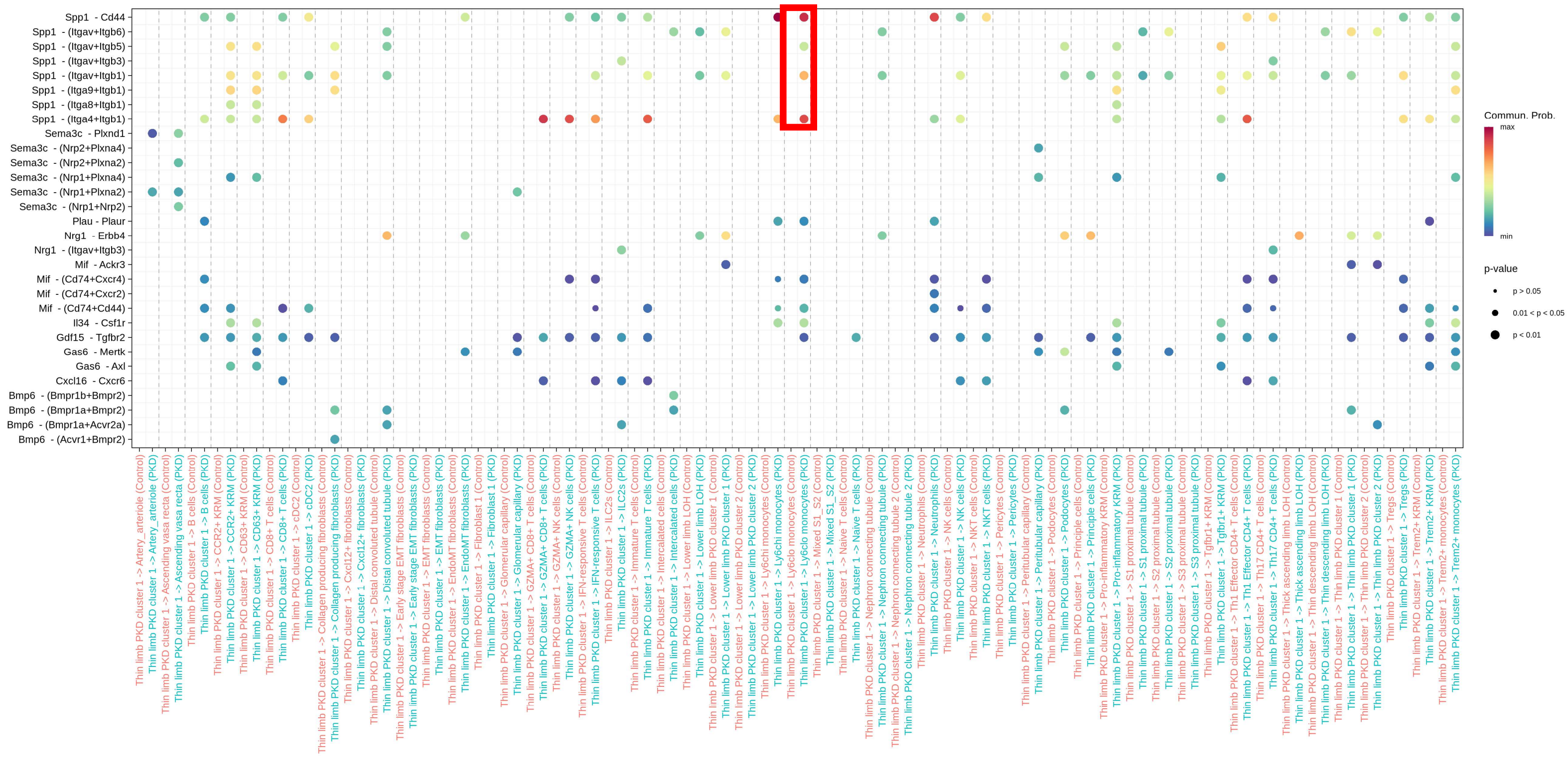

Supplemental figure 30

#### Lower limb PKD cluster 2

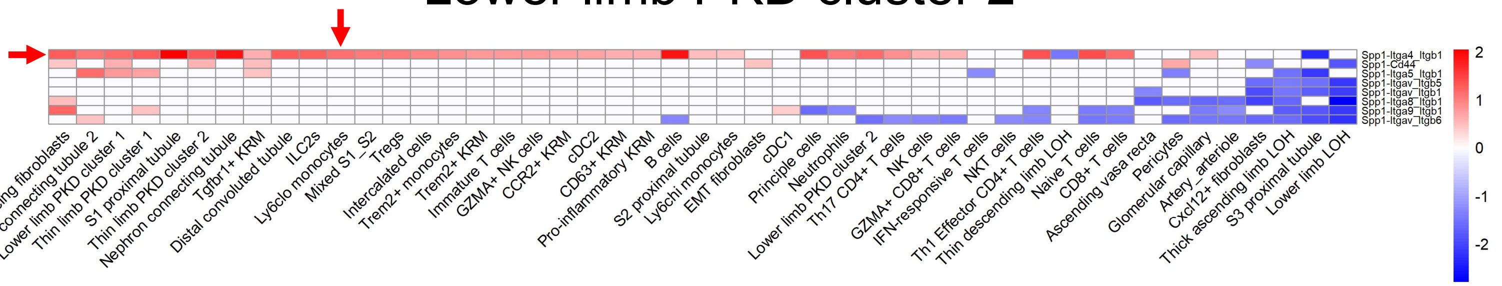


#### Thin limb PKD cluster 1

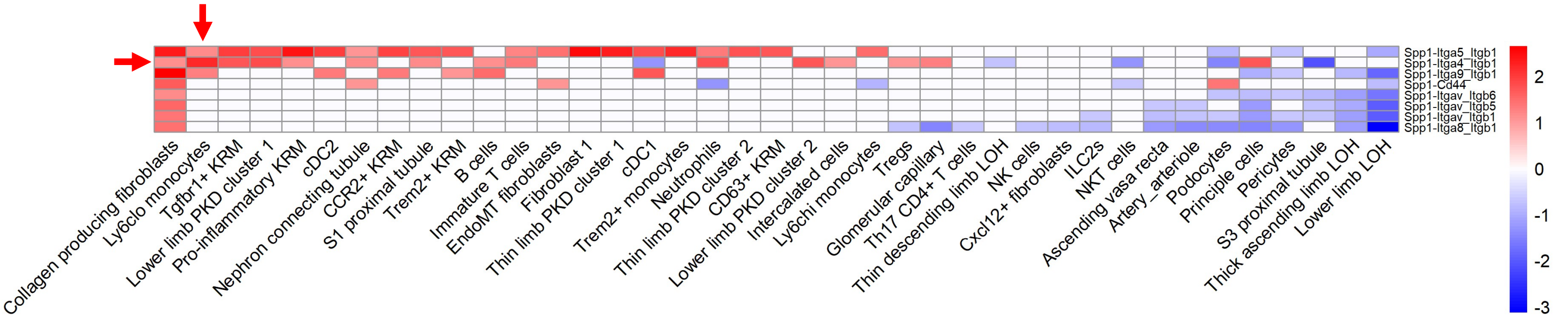

### Supplemental figure 31

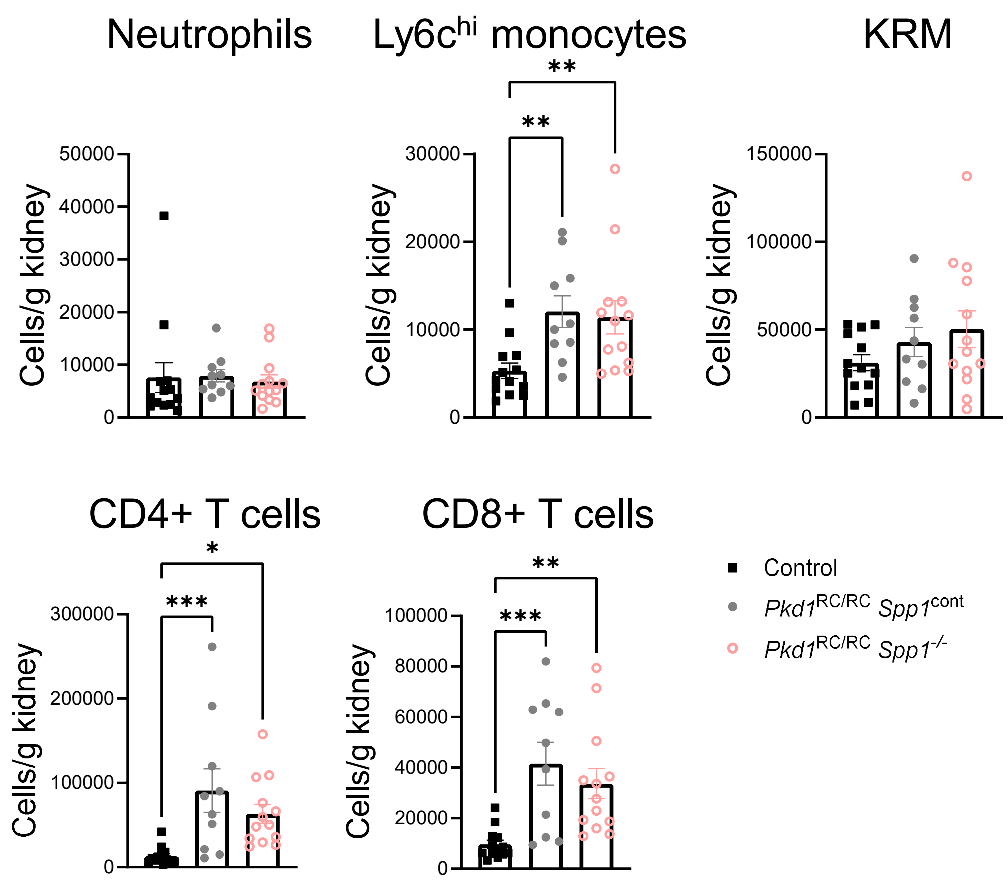

Supplemental figure 32

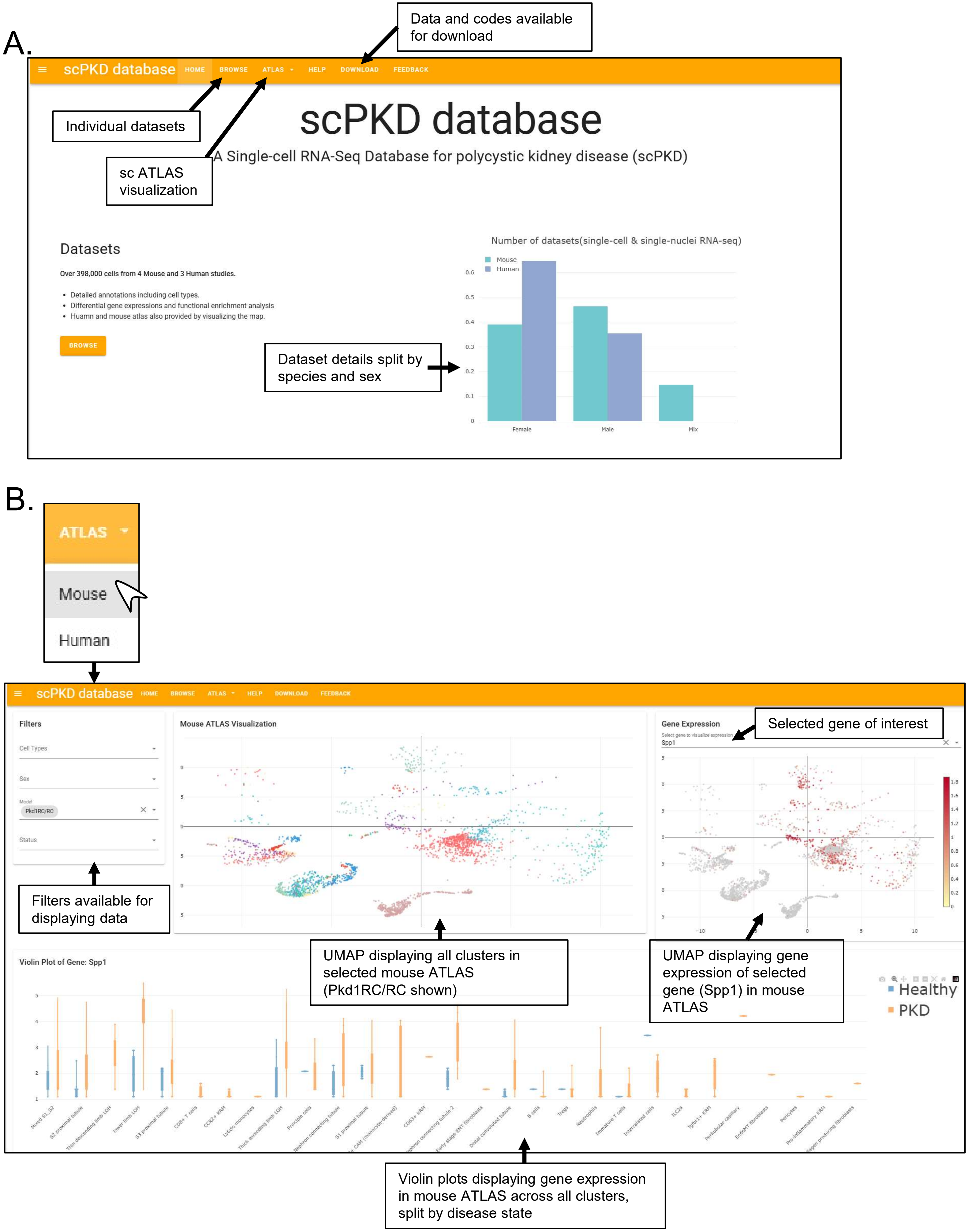
