## Supplemental Methods for "A cross model spatial and single-cell atlas reveals the conserved involvement of osteopontin in polycystic kidney disease"

*IACUC approval:* All animals were maintained in Association for Assessment and Accreditation of Laboratory Animal Care (AALAC) International-accredited facilities in accordance with Institutional Animal Care and Use Committee regulations at the University of Oklahoma Health Sciences Center under approved protocol number 23-002-CHIX.

*Mouse models used for scRNAseq data:*

*Ift88:* Seven to nine-week-old female C57BL/6J CAGGCre^ERT2^ *Ift*88^f/f^ mice received a single intraperitoneal (IP) injection of tamoxifen at 6 mg/40g body weight for 3 consecutive days, as previously done by our lab^1, 2^. CAGGCre^ERT2^ *Ift*88^f/f^ mice were harvested ~6-7 months post tamoxifen induction and kidneys isolated for scRNAseq as described below. As scRNAseq is a discovery-based approach and not meant for rigid quantitative analysis, scRNAseq experiments were done on N=2 mice per group (2 control, 2 cystic; all female). Raw data are available in GEO (GSE193528)^1^.

*Pkd2:* Seven to nine-week-old male and female C57BL/6J CAGGCre^ERT2^ *Pkd2*^f/f^ mice received a single intraperitoneal (IP) injection of tamoxifen at 6 mg/40g body weight for 3 consecutive days. CAGGCre^ERT2^ *Pkd2*^f/f^ mice were harvested ~4-5 months post tamoxifen induction and kidneys isolated for scRNAseq as described below. scRNAseq experiments were done on N=2 mice per group (2 control, 2 cystic; all female). Raw data are available in GEO (GSE297737).

Injured *Ift88:* Seven to nine-week-old male and female C57BL/6J CAGGCre^ERT2^ *Ift*88^f/f^ mice received a single intraperitoneal (IP) injection of tamoxifen at 6 mg/40g body weight for 3 consecutive days. Three weeks post tamoxifen injection, mice were subjected to 30 minutes of sham surgery or unilateral ischemia reperfusion injury as described in our previous manuscript that generated the scRNAseq data^1^. scRNAseq experiments were done on N=2 mice per group (2 control injured, 2 cystic sham operated, 2 cystic injured; all female). Raw data are available in GEO (GSE193528)^1^.

*Pax8* teto^cre^ *Pkd1*^f/f^: Pax8 mice received doxycycline from P27 to P56 followed by harvest of kidneys at P66, P100, P130 as described^3^. scRNAseq experiments were done on N=2-3 mice per group (P66: 2 control, 2 cystic; P100: 3 control, 3 cystic; P130: 3 control, 3 cystic; all male). Raw scRNAseq data was obtained from GEO (GSE268494)^3^.

*Pkhd1*^cre^ *Pkd1*^f/f^: Kidneys were harvested at P7, P14, P21 as previously described^4^. scRNAseq experiments were done on 2-3 mice per group (male and female mix). Cellranger outputs were obtained from originating authors. No GEO data is available for these samples.

*Pkd1*^RC/RC^: Control (*Pkd1*^RC/WT^) and cystic (*Pkd1*^RC/RC^) kidneys were harvested at one year of age and processed for scRNAseq as described below. scRNAseq experiments were done on N=2-3 mice per group (2 control, 3 cystic; 1M control, 1 F control; 1M cystic, 2F cystic). Raw data are available in GEO (GSE298379).

*Sample prep for single cell RNA sequencing:* For in house scRNAseq experiments (*Pkd1*^RC/RC^, *Pkd2*), kidneys were minced and digested in 1ml of 2.5 mg/ml type II collagenase (Worthington, cat No; LS004176), 7.5 mg/ml B. licheniformis cold activated protease (Creative enzymes, Cat no: NATE-0633), and 125 U/ml DNase (Sigma-Aldrich, Cat no: D5025) in D-PBS for ~45 minutes at 12C as previously described^1^. After digestion, kidney tissue was passed through individual 70-µm cell-strainers yielding single-cell suspensions. Cells were centrifuged at 300 X g for 5 minutes at 4ºC, resuspended in ACK red blood cell lysis buffer, and incubated at 37ºC for 5 minutes. After 5 minutes, 10 ml RPMI was added to each tube, cells were spun for 5 minutes at 4C, and cells were resuspended in 0.04% BSA with FC blocking solution (1:200, BioXcell; catalog #: BE0307) for 30 minutes on ice. After blocking, cells were spun and incubated with the following antibodies in 0.04% BSA for 30 minutes: PE rat anti-mouse CD45 (catalog no. 12–0451, 30-F11; eBioscience), and Fixable Aqua Dead Cell Stain (catalog no. L34957; Invitrogen). Stained cells were then washed with 0.04% BSA and resuspended in 0.04% BSA for sorting. We sorted ~25,000 live, CD45+ or CD45- cells into individual BSA coated tubes. Cells were counted and approximately 5,000 cells from each group were combined into a single tube followed by performing bead emulsion and 10X genomics as described below.

*10X genomics*: 10x Chromium single cell libraries were prepared according to the standard protocol outlined in the manual. Briefly, sorted single cell suspension, 10x barcoded gel beads, and oil were loaded into Chromium™ Single Cell Chip G to capture single cells in nanoliter-scale oil droplets by Chromium™ Controller and to generate Gel Bead-In-EMulsions (GEMs). We used “Chromium™ Single Cell 3' GEM, Library & Gel Bead Kit v3.1” and chip G for these experiments. Full length cDNA libraries were prepared by incubation of GEMs in a thermocycler machine. GEMs containing cDNAs were broken and all single cell cDNA libraries were pooled together, cleaned using DynaBeads MyOne™ Silane beads (Fisher PN 37002D), and pre-amplified by PCR to generate sufficient mass for sequencing library construction. Sequencing libraries were constructed by following the steps: cDNA fragmentation, end repair & A-tailing, size selection by SPRIselect beads (Beckman Coulter, PN B23318), adaptor ligation, sample index PCR amplification, and a repeat of SPRIselect beads size selection. The final constructed single cell libraries were sequenced by Illumina Nextseq2000 machine with total reads per cell targeted for a minimum of 25,000.

*Single cell sequencing data processing*: The 10X Genomics Cellranger software (version 7.1.0), ‘mkfastq’, was used to create the fastq files from the sequencer. Following fastq file generation, Cellranger ‘count’ was used to align the raw sequence reads to the reference genome using STAR. The ‘count’ software created 3 data files (barcodes.tsv, features.tsv, matrix.mtx) that were loaded into the R package Seurat version 5.2.0 ^5^, which allows for selection and filtration of cells based on QC metrics, data normalization and scaling, and detection of highly variable genes. Next, we used DoubletFinder^6^ and Soupx^7^ to remove doublets and ambient RNA from the data prior to data integration. We followed the Seurat vignette (<https://satijalab.org/seurat/pbmc3k_tutorial.html>) to create the Seurat data matrix object for each individual object followed by combining individual replicates together using the ‘merge’ function. To generate the integrated scRNAseq atlas shown in Figure 1B, we used SCT-RPCA on the layered Seurat object. After subsetting stromal cells, we used SCT-Harmony and the previous RPCA-integration. After all integration and merging steps, we removed low quality cells by keeping all genes expressed in greater than 3 cells and cells with at least 200 detected copies. Cells with mitochondrial gene percentages over 50% and unique gene counts greater than 3,000 or less than 200 were discarded, as previously established in the kidney^1^. The data was normalized using Seurat’s ‘NormalizeData’ function, which uses a global-scaling normalization method, LogNormalize, to normalize the gene expression measurements for each cell to the total gene expression. Highly variable genes were then identified using the function ‘FindVariableGenes’ in Seurat. We also regressed out the variation arising from library size and percentage of mitochondrial genes using the function ‘ScaleData’ in Seurat. We performed principal component analysis (PCA) of the variable genes as input and determined significant PCs based on the ‘JackStraw’ function in Seurat. The first 10 PCs were selected as input for Uniform Manifold Approximation and Projection (UMAP) dimensionality reduction using the functions ‘FindClusters’ and ‘DimPlot’ in Seurat. To identify differentially expressed genes in each cell cluster, we used the function ‘FindAllMarkers’ in Seurat on the normalized gene expression data. Detailed code for generating and analyzing scRNAseq data can be found in github as outlined in the “code availability” section below.

*DecoupleR:* DecoupleR^8^ for pathway and transcription factor inference was done using the integrated Seurat object and the standard vignette available on github (<https://saezlab.github.io/decoupleR/articles/pw_sc.html>).

*Differential expression testing using pseudobulked data:* For the differential expression testing, raw counts for each cluster for each experimental condition were pseudobulked using the ‘AggregrateExpression’ function and the counts slot. Differential gene expression testing was done using DESeq2 and plotted using pheatmap. Due to the high variance across groups, we used both adjusted p value (<0.05) and non-adjusted p value (<0.05) when plotting differentially expressed genes.

*CellChat analysis for cell-cell communication in scRNAseq data:* CellChat^9^ was performed on the fully annotated scRNAseq atlas as shown in Supplemental figure 9B using the standard vignette for analyzing multiple scRNAseq samples (<https://htmlpreview.github.io/?https://github.com/jinworks/CellChat/blob/master/tutorial/Comparison_analysis_of_multiple_datasets.html>). Briefly, we created separate CellChat objects for control and PKD samples using the group metadata column followed by merging the two CellChat objects into a single object using the ‘mergeCellChat’ function. We then calculated the netcentrality scores using the ‘netAnalysis_computeCentrality’ function followed by performing downstream analysis using the standard vignette.

*10X Visium* (*Ift88*): Paraffin embedded kidney tissue from *Ift88* mice were used for spatial transcriptomics (2 controls, 2 cystic; all mice female; 4 total). Initially, we sectioned 4-5 20 μm sections into an Eppendorf tube, isolated RNA using the RNeasy FFPE kit (Qiagen, Catalog number 73504) and analyzed the quality of the isolated RNA using an Agilent Bioanalyzer. We ensured that the DV200 for all samples was greater than 45% prior to performing 10X Visium. For sectioning in preparation of Visium, paraffin embedded blocks were chilled on ice prior to mounting onto the active sequencing areas (6 mm × 6 mm) of the 10X Genomics Visium slides. The slides were deparaffinized and H&E stained to perform high quality imaging on a Nikon TE2000-E microscope. After imaging, samples were processed using the 10x Genomics Visium Spatial for FFPE Gene Expression Kit, mouse 6.5mm and their established protocols. Decrosslinking was performed on slide to release the RNA. Probe Hybridization was immediately performed on the slides followed directly by probe ligation, probe release, and extension with UMIs and spatial barcodes. Products were eluted from the slide for library preparation and indexing. Library quality was assessed using Agilent’s Bioanalyzer and Invitrogen’s Qubit 4 Fluorometer. Sequencing depth for each sample was estimated based on the approximate Capture Area covered (%) by the tissue section. Purified libraries were normalized, pooled, and sequenced on Illumina’s NovaSeq platform, targeting 125 million reads per sample using dual indexing. Resulting FASTQ files were aligned to mm10 reference, manually aligned to respective hematoxylin and eosin stained sections and normalized using 10X Genomics Space Ranger count (spatial 3′ v1; spaceranger-1.2.1). Each of the sequenced libraries covered approximately 1,500-2,000 barcoded spots (Control *Ift88* 1- 1935 spots, Control *Ift88* 2- 1533 spots, *Ift88* 1- 2194 spots, *Ift88* 2- 1945 spots) across the embedded capture probe area. Median genes per spot ranged from 5218 to 7278 with mean reads per spot ranging from 37,845-92,348.

*10X Visium* (*Pkd1*^RC/RC^): Fresh frozen kidney tissue from ~7-month-old *Pkd1*^WT/WT^ and *Pkd1*^RC/RC^ mice were used for spatial transcriptomics (2 controls (1M, 1F), 2 cystic (1M, 1F); 4 total). Initially, we sectioned 4-5 20 μm sections into an Eppendorf tube, isolated RNA using the RNeasy kit (Qiagen, Catalog number 74104) and analyzed the quality of the isolated RNA using an Agilent Bioanalyzer. We ensured that the DV200 for all samples was greater than 45% prior to performing 10X Visium. 10x Visium slides with fresh frozen kidney tissue were H&E stained followed by imaging using a Nikon TE2000-E microscope. After imaging, samples were processed using the 10x Genomics Visium Spatial Gene Expression Kit, mouse 6.5mm and their established protocols. Permeabilization and reverse transcription were performed followed by second strand synthesis and denaturation for sample removal from the slide. Off the slide, cDNA amplification and QC were performed before standard library construction. Library quality was assessed using Agilent’s Bioanalyzer and Invitrogen’s Qubit 4 Fluorometer. Sequencing depth for each sample was estimated based on the approximate Capture Area covered (%) by the tissue section. Purified libraries were normalized, pooled, and sequenced on Illumina’s NovaSeq platform, targeting 125M reads per sample using dual indexing. Resulting FASTQ files were aligned to mm10 reference, manually aligned to respective hematoxylin and eosin stained sections and normalized using 10X Genomics Space Ranger count (spatial 3′ v1; spaceranger-1.2.1). Each of the sequenced libraries covered approximately 1,000-2,000 barcoded spots (Control *Pkd1*^WT/WT^- 2289 spots, Control *Pkd1*^WT/WT^- 1729 spots, *Pkd1*^RC/RC^- 2289 spots, *Pkd1*^RC/RC^- 874 spots) across the embedded capture probe area. Median genes per spot ranged from 4230 to 4664 with median reads per spot ranging from 40,815-54,463.

*Processing of spatial transcriptomics data.* Initial processing of spatial transcriptomics data was done using scanpy and the standard vignette available on their website (<https://scanpy-tutorials.readthedocs.io/en/latest/spatial/basic-analysis.html>). The resulting adata objects were used for downstream deconvolution using TACCO.

*Cell type deconvolution using transfer of annotations to cells and their combinations (TACCO).* To deconvolute spots into predicted cell types, we used TACCO^10^, a computational framework for the decomposition and annotation of diverse biological measurements in single cell and spatial genomics and our integrated, annotated scRNAseq atlas shown in Figure 11. Default TACCO settings were applied and the output provides a compositional annotation of spatial spots (<https://github.com/simonwm/tacco>). Cell type deconvolutions for control and PKD samples are shown in Supplemental figures 14-21.

*Analysis of cell-cell communication in spatial data using collective optimal transport.* For analysis of intercellular signaling networks in spatial data, we used collective optimal transport (COMMOT)^11^, a tool for analyzing intercellular communication networks using single-cell RNA-sequencing data. This vignette (<https://commot.readthedocs.io/en/latest/index.html>) uses collective optimal transport to construct CCC networks for the ligand-receptor pairs with a spatial distance constraint of 200μm (coupling between cells with distance greater than 200 μm is prohibited). Calculated CCC scores for each ligand receptor pair between each cell type in the scRNAseq atlas were extracted from python and imported in R for visualization.

*Identification of cystic niches:* To identify intercellular signaling interactions (Ligand-Receptor pairs) within cystic niches, we outlined cystic regions using ImageJ software. Next, we extracted the spatial coordinates associated with outlined cystic regions using the metadata associated with each .tif file, imported the spatial coordinates of cystic regions into python, and identified spots touching cystic regions in each sample. The outlined regions and selected cystic spots for each sample are shown in Figure 7D. To infer signaling within cystic niches, we then selected all spots that were within 1 spot (~55μm) of the outlined region (Supplemental figure 23A). Next, we extracted CCC scores from COMMOT from each of cystic and non-cystic niches and plotted the top 50 ligand-receptor interactions between cell types in Supplemental figure 23 B,C.

*Analysis of variable genes across cyst size:* To understand how cyst size impacts gene expression, we quantified the area within each outlined cystic region in python followed by binning cysts into 3 groups (large, medium, small) across all four cystic replicates (Supplemental figure 24A). We then extracted gene expression from each spot found within each of the three different sized cysts followed by identification of the top 100 highly variable genes (HVGs), which is shown as a principle component plot in Supplemental figure 24B. We also analyzed genes that were differentially expressed between different sized cysts in relation to one another and non-cystic regions using DESeq2 (Supplemental figure 24C, 25). For individual cyst analysis, we identified the top 100 HVGs at the spot or cyst level and plotted the results as a heatmap (Supplemental figure 26).

*Data availability for 10X Visium data:* Raw sequencing data for spatial transcriptomics studies (both *Pkd1*^RC/RC^ and *Ift88*) are available in GEO (GSE299863).

*Pkd1*^RC/RC^ *Spp1 knockout mice: Pkd1*^RC/RC^ *Spp1*^cont^ or *Pkd1*^RC/RC^ *Spp1*-/- mice were harvested at ~1 year of age. Sample size was calculated based on initial data assuming a 25% difference in cystic index, 20% deviation, α = 0.05, and β = 0.2. The power calculation indicated a minimum of N=10 mice per group. Outliers were identified as having a z score greater than 3 using the outlier test. For the manuscript, one mouse in the *Pkd1*^RC/RC^ *Spp1*^cont^ and one mouse in the *Pkd1*^RC/RC^ *Spp1*-/- group was excluded as it had a z score greater than 3.

*Tissue processing.* *Pkd1*^RC/RC^ *Spp1* mice were perfused with PBS to remove contaminating immune cells in the vasculature. The left kidney was removed and put into RPMI 1640 on ice while the right kidney was cut in half and specimens were immediately immersed in in 10%(wt/vol) NBF overnight at 4ºC. The next day, samples were either switched to 30% (wt/vol) sucrose overnight at 4 ºC for cryopreservation or 70% ethanol for paraffin embedding. Paraffin embedded tissue was sectioned into 5 μm slices for use in downstream applications.

*Quantification of cystic severity*: 5-μm sections were cut from paraffin-embedded kidneys and stained with hematoxylin-eosin (H&E). To identify cysts, we began by quantifying the average diameter and standard deviation of a normal tubule using control, Cre negative mice. Cysts were identified as any opening (white space) that was greater than 3 standard deviations above the average diameter of a normal tubule. To ensure that we did not include blood vessels and other artifacts that were the result of tissue processing in our cystic calculations, we went back and individually analyzed all particles that were identified as cysts using ImageJ. After removal of non-cystic structures, we calculated cystic index as the total cystic area divided by the total area of the sectioned kidney. This quantification was done using at least ½ of the kidney for all animals. A technician blinded to the treatment modality performed the quantification and analysis.

*Flow cytometry:* Flow cytometry experiments were performed following published protocols^12^. Briefly, animals were perfused with PBS before the left kidney was extracted and minced followed by enzymatic digestion for 30 minutes at 37 C. Digestion buffer consisted of 1mg/mL collagenase Type I (Sigma-Aldrich, Catalog#: C0130-100MG) and 100 U/ml DNase I (Sigma-Aldrich, Catalog#: D5025-15KU). Cells were filtered through a 70μm strainer (MidSci, Catalog#: 70ICS), red blood cells were lysed (ACK lysis buffer, Quality Biological, Catalog#: 10128-802) and cells were resuspended in 1% BSA in PBS containing Fc blocking solution. After 30 minutes on ice, 2x10^6^ cells were stained with the appropriate amount of primary antibody (Supplemental table 5). Following a 30-minute incubation at room temperature, cells are washed and fixed with 2% PFA on ice for 30 minutes. Lastly, cells are resuspended in PBS before being analyzed on a Cytek Aurora. All analysis were conducted using FlowJo V10.9.0 software.

*Blood Urea Nitrogen (BUN) analysis.* Blood was collected from mice prior to perfusion with PBS via cardiac puncture and placed in EDTA coated tubes (Monoject, Catalog number: 8881311743). Blood samples were allowed to rock at room temperature for ~30 minutes prior to processing. Collected blood was centrifuged for 10 minutes at 1200 × g and the supernatant was transferred to an Eppendorf tube and stored at −80°C. Resulting plasma was analyzed for changes in BUN using the QuantiChrom Urea Assay Kit (DIUR-100; BioAssay Systems) according to the manufacturer’s instructions.

*Code availability:* All code for these experiments can be found in out labs github account (kzimmer1) under the repository name “Mouse-SingleCell-Atlas”.

*Blinding and randomization*: All mice used for scRNAseq were randomized prior to treatments. On the day of harvest, the flow cytometry and scRNAseq was performed by a scientist who was blinded to experimental groups. Bead capture, cDNA library preparation, sequencing, file conversion, and FASTq file generation were performed by the flow cytometry core at UAB or OMRF in a blinded manner. Data were processed in R in a blinded manner until the cluster composition and cell numbers were revealed.

*Statistics.* To determine statistical significance between experimental groups, we performed a one-way or two-way ANOVA without correction for multiple comparisons. For head-to-head comparisons, we performed a Student’s T test. Residual plots were used to determine normality of the data. Data was considered statistically significant for P values less than 0.05.
